# Global identification of direct SWI/SNF targets reveals compensation by EP400

**DOI:** 10.1101/2023.03.07.531379

**Authors:** Benjamin J.E. Martin, Eileen F. Ablondi, Christine Goglia, Karen Adelman

## Abstract

Mammalian SWI/SNF chromatin remodeling complexes move and evict nucleosomes at gene promoters and enhancers to modulate DNA access. Although SWI/SNF subunits are commonly mutated in disease, therapeutic options are limited by our inability to predict SWI/SNF gene targets and conflicting studies on functional significance. Here, we leverage a fast-acting inhibitor of SWI/SNF remodeling to elucidate direct targets and effects of SWI/SNF. Blocking SWI/SNF activity causes a rapid and global loss of chromatin accessibility and transcription. Whereas repression persists at most enhancers, we uncover a compensatory role for the EP400/TIP60 remodeler, which reestablishes accessibility at most promoters during prolonged loss of SWI/SNF. Indeed, we observe synthetic lethality between EP400 and SWI/SNF in lung cancer cell lines and human cancer patient data. Our data define a set of molecular genomic features that accurately predict gene sensitivity to SWI/SNF inhibition in diverse cancer cell lines, thereby improving the therapeutic potential of SWI/SNF inhibitors.

**Highlights:**

- Genes repressed by long-term SWI/SNF loss do not accurately reflect direct targets
- Promoters that fail to reestablish activity are those lacking a compensatory remodeler
- Accessibility and activity recovery requires EP400, synthetically lethal with SWI/SNF
- SWI/SNF dependence in cancer cells can be predicted from the promoter chromatin state

## INTRODUCTION

Gene activation requires that transcription factors (TFs) and the transcription machinery can access DNA, both at gene promoters and at cis-regulatory enhancers.^1, 2^ DNA accessibility is generated by chromatin remodelers such as the mammalian SWI/SNF complexes, which use energy from ATP to slide nucleosomes or evict them from DNA. These actions create nucleosome-depleted regions at promoters and enhancers that facilitate TF binding and transcription initiation.^3, 4^ Further, SWI/SNF has been implicated in rendering chromatin more dynamic to help RNA polymerase II (Pol II) overcome nucleosome barriers within gene bodies.^5^ Accordingly, chromatin remodeling by SWI/SNF is critical to the establishment of appropriate gene expression patterns.^6–8^

Emphasizing the crucial role of SWI/SNF, the complex is mutated in >20% of cancers, with SWI/SNF subunits frequently found to contain driver mutations.^9, 10^ However, a comprehensive understanding of the targets and cellular consequences of SWI/SNF activity has remained elusive, as RNAi-mediated depletion, genomic knockout, and mutational studies have reported varied, often conflicting, conclusions about the function of chromatin remodeling by SWI/SNF.^11–13^ Specifically, current models for SWI/SNF function range from inhibition of enhancer transcription,^11, 14^ to repression of specific gene sets,^3, 15, 16^ to activation of specific enhancers.^7, 17, 18^ These discrepancies likely result from the extended time required to sufficiently deplete SWI/SNF proteins with these strategies, such that the direct effects of SWI/SNF loss are obscured by indirect effects and compensatory mechanisms.

For these reasons, the development of fast-acting and specific inhibitors and degraders of the paralogous SWI/SNF ATPase subunits BRG1 and BRM represent valuable tools towards elucidating the direct role of SWI/SNF-mediated nucleosome remodeling in living cells.^19^ Indeed, treatment of mouse and human cells with fast acting BRG1/BRM inhibitors BRM011 and BRM014 markedly reduced chromatin accessibility at many regulatory loci within minutes, indicating that the maintenance of open chromatin at these sites is dependent upon continuous catalytic activity of SWI/SNF.^20, 21^ These results are consistent with previous work in S. cerevisiae demonstrating that constant chromatin remodeling is required to maintain appropriate genomic accessibility patterns.^22–24^ Importantly, the effects observed upon treatment of cells with BRM014 were highly similar to those obtained when BRG1 was subjected to targeted protein degradation, validating inhibitor specificity.^20, 25^ Further, the development of SWI/SNF mutants with resistance to inhibitor compounds selectively identified mutated residues located within the catalytic active site of the BRM/BRG1 ATPases.^26^

Surprisingly, despite broad reduction in enhancer accessibility and transcription factor occupancy observed when BRG1/BRM were inhibited or degraded, this resulted in limited and highly selective effects on gene expression.^20, 21, 25^ These findings raised critical questions about the functional relevance of SWI/SNF mediated remodeling at regulatory elements.

One possibility suggested by the recent data is that alternate mechanisms exist to allow a majority of genes to maintain expression in the absence of BRG1/BRM activity. To address this possibility, and to identify potential compensatory chromatin remodelers, we probed the direct impact of SWI/SNF inhibition on enhancer and gene activity using time-resolved assays of chromatin accessibility and active transcription in mouse embryonic stem cells (mESCs). We find that SWI/SNF is globally and continuously required for chromatin accessibility and transcription initiation at both enhancers and promoters. However, whereas enhancers are persistently repressed during SWI/SNF inhibition, many promoters recover accessibility and transcription activity. Our analyses reveal that promoters that fail to recover are characterized by weak chromatin accessibility and an enrichment of H3K4 monomethylation (H3K4me1) over trimethylation (H3K4me3). Importantly, these chromatin features defined in mESCs can predict gene sensitivity to SWI/SNF perturbation in non-small cell lung cancer (NSCLC) and prostate cancer cell lines. Thus, our work establishes a prognostic framework for identifying genes that will be sensitive to SWI/SNF loss or inhibition in disease contexts. Further, we demonstrate that the compensation for loss of SWI/SNF activity is mediated by the EP400/TIP60 coactivator complex, which interacts with and is recruited by H3K4me3. Accordingly, EP400 gains increased importance in cells wherein SWI/SNF is perturbed and EP400 loss greatly sensitizes cells to loss of SWI/SNF activity.

## RESULTS

### SWI/SNF inhibition causes widespread reduction in enhancer activity

To characterize changes in chromatin accessibility upon SWI/SNF inhibition in mESCs, we began by systematically identifying active promoters and enhancers genome-wide (Figure 1A). ATAC-seq data from untreated mESCs was used to define a set of peaks corresponding to regions of accessible chromatin (N=83,201). We then used PRO-seq, which captures nascent RNA associated with engaged RNAPII,^27^ to define sites of active transcription at both annotated and unannotated RNA loci. Approximately 20% of ATAC-seq peaks were located within 1.5 kb of an active annotated transcription start site (TSS), with a median distance of 112 bp between these peak centers and the nearest active TSS (Figure S1A, see STAR methods). These peaks were therefore designated as “promoter peaks” and, to facilitate subsequent analysis, were centered on the active TSS (N=13,536; Figure S1B). As synthesis of enhancer RNA (eRNA) is a sensitive hallmark of active enhancers,^28–30^ promoter-distal ATAC-seq peaks with associated PRO-seq signal^31^ were classified as putative enhancers (N=32,149). Consistent with this designation, transcribed ATAC-seq peaks were enriched for acetylated histone H3K27 as compared to non-transcribed peaks (Figure S1C) and showed the enrichment of H3K4 monomethylation (H3K4me1) over H3K4 trimethylation (H3K4me3) that is often considered a hallmark of enhancers (Figure S1D). ChIP-seq for BRG1, the SWI/SNF ATPase subunit expressed in mESCs,^32^ demonstrated BRG1 occupancy at both promoters and enhancers (Figure 1A), consistent with a broad role for SWI/SNF in chromatin remodeling.^2, 33^

**Figure 1.**
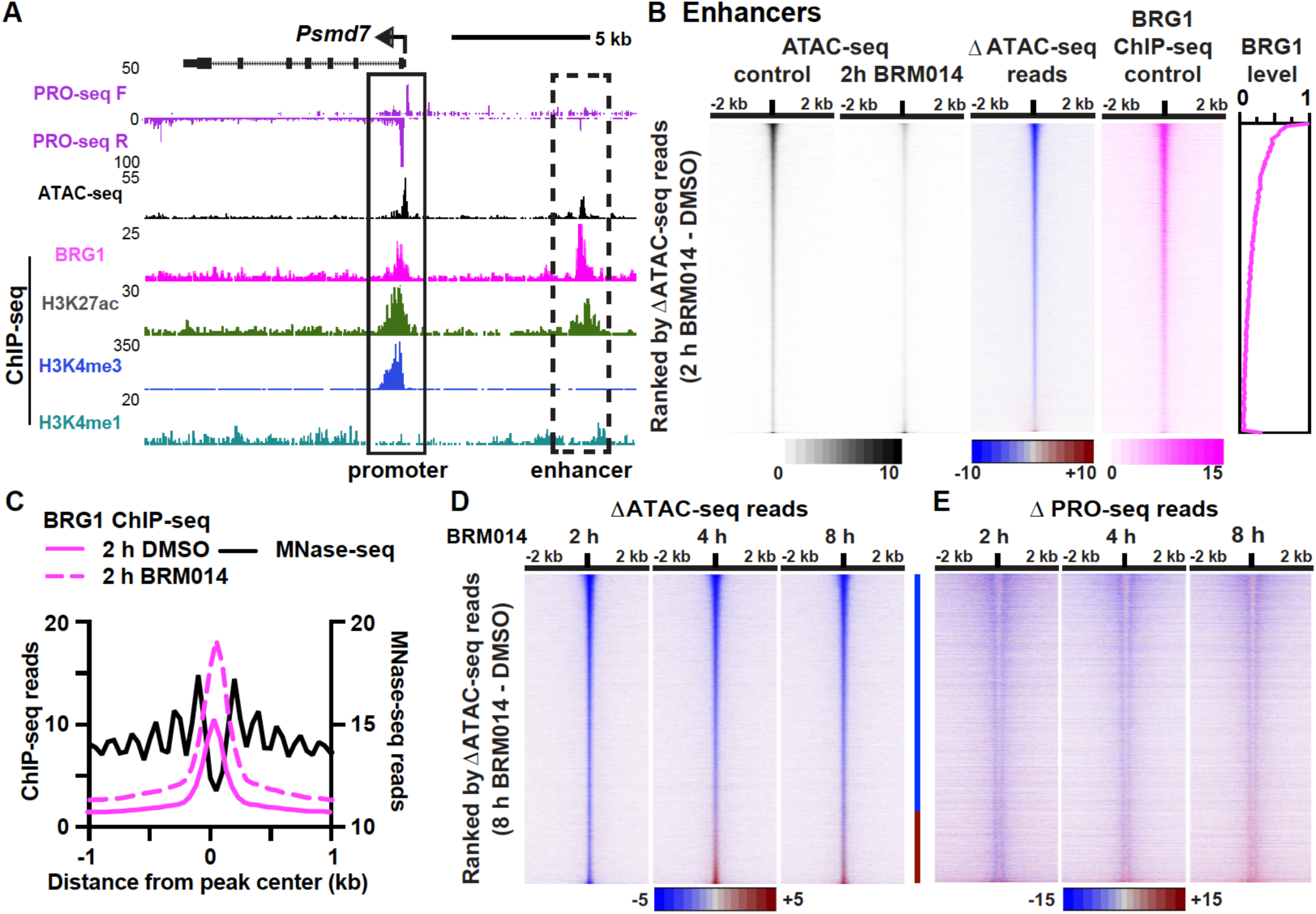
Enhancer accessibility and activity require SWI/SNF activity. (**A**) Genome browser view of the *Psmd7* promoter (solid box) and associated enhancer (dashed box) with PRO-seq, ATAC-seq, BRG1 ChIP-seq, H3K27ac ChIP-seq,^60^ H3K4me3 ChIP-seq^60^ and H3K4me1 ChIP-seq data.^61^ **(B)** Heatmap representation of the effects of 2 h BRM014 treatment on ATAC-seq signal at enhancers (n = 32,149). Normalized data from combined replicates (n = 3 per condition) were aligned to the enhancer center. Sites are ranked by difference in ATAC-seq reads (Enhancer center +/- 300 bp) between 2 h BRM014 and 2 h DMSO control. BRG1 ChIP-seq signal is shown in the same rank order, as is relative BRG1 signal (summed +/- 500 bp from peak centers). **(C)** Aggregate plot of quantitative BRG1 ChIP-seq signal (n = 2 per condition) at enhancers from 2 h DMSO- and BRM014-treated cells. Average MNase-seq^28^ profiles are shown to indicate the position of the NDR. Data are graphed in 50 bp bins. **(D and E)** Difference in ATAC-seq signal (D) and PRO-seq signal (E) after BRM014 treatment (n ≥ 2 per condition) for all enhancers. Data were aligned to the enhancer center and rank ordered by the difference in enhancer ATAC-seq reads after 8 h treatment. Blue line between heatmaps indicates the 77% of enhancers that fail to recover accessibility, while red line indicates enhancers that regain accessibility. See also Figures S1 and S2.

We then treated mESCs with BRM014 (at 1 µM) or DMSO for 2 h and performed ATAC-seq, with *Drosophila* spike in controls to allow for accurate quantification. These data demonstrated that inhibition of SWI/SNF activity broadly reduces chromatin accessibility at enhancers (Figure 1B, >98% of enhancers affected). This finding is consistent with prior work in mESCs which demonstrated strongly reduced accessibility and occupancy of transcription factors (TFs) at regulatory loci following SWI/SNF loss, and clarifies that this loss of accessibility occurs at nearly all enhancer loci.^8, 20, 21^ Supporting that these rapid consequences of SWI/SNF inhibition represent direct effects, the observed magnitude of accessibility changes at enhancers agrees well with BRG1 occupancy, such that the enhancers most strongly affected by SWI/SNF inhibition are those most highly bound by the complex (Figure 1B, BRG1 ChIP-seq). Investigation of BRG1 binding to chromatin using quantitative ChIP-seq after 2h BRM014 treatment demonstrated an increase in BRG1 occupancy within the enhancer peak, centered over the nucleosome-depleted region (NDR) (Figure 1C). This augmented BRG1 occupancy is consistent with biochemical experiments indicating that blocking ATP hydrolysis slows SWI/SNF release from chromatin.^34^ Short term inhibition of BRG1 thus does not displace SWI/SNF from enhancers, as would protein depletion or degradation. Consequently, this system provides mechanistic insights into the role of the BRG1 ATPase under conditions wherein the SWI/SNF complex remains properly localized. Western blotting confirmed that treatment with BRM014 for up to 24 h had no detectable effect on levels of SWI/SNF subunits (Figure S1E).

We next determined chromatin accessibility following extended inhibition of SWI/SNF activity, focusing on BRM014 treatment for 4 and 8 h, time points at which we observed no defects in cell proliferation or morphology (Figures S1F and S1G). Indicative of a continued dependence of enhancers on SWI/SNF for maintenance of open chromatin, accessibility was reduced at 77% of enhancers throughout an 8 h treatment with BRM014 (Figure 1D, indicated by blue line at right). Given this striking loss of accessibility during prolonged inhibition of SWI/SNF, and recent reports that BRM014 markedly reduces the occupancy of key TFs at enhancers, one might predict that enhancer RNA synthesis would also be repressed by BRM014. Accordingly, analysis of nascent RNA synthesis at enhancers using PRO-seq demonstrates a broad reduction in eRNA transcription across the BRM014 treatment time course (Figure 1E). These results contrast with earlier suggestions that SWI/SNF suppresses enhancer transcription based on long-term depletion approaches,^11^ but are consistent with recent work using acute SWI/SNF perturbation.^20, 21, 25^ We conclude that chromatin remodeling by SWI/SNF is necessary for the sustained activity of most enhancers.

Notably, enhancers that were sensitive to loss of SWI/SNF activity across the 8 h time course (Figure 1D, indicated by blue line) were enriched in binding of the pluripotency-associated transcription factors OCT4, SOX2, and NANOG (OSN) as compared to enhancers that recovered accessibility (Figures S2A-S2C). This finding emphasizes that the presence of TFs considered to be pioneer factors does not render an enhancer less dependent on chromatin remodelers.^8, 21^ By contrast, CTCF occupancy was enriched at the subset of enhancers that recovered accessibility following SWI/SNF inhibition (Figures S2A, S2D and S2E).^4, 21^ This observation suggests that the CTCF-associated chromatin remodeler SNF2H (SMARCA5)^35, 36^ might serve to maintain open chromatin at these sites during SWI/SNF inhibition. Consistent with this idea, we found SNF2H enrichment at enhancers that retained accessibility following BRM014 treatment (Figure S2E). Moreover, accessibility at this subset of enhancers was sensitive to knockout of SNF2H (Figure S2F), confirming a role for remodeling by SNF2H at these loci. Together, these data indicate that most enhancers in mESCs, including those bound by pioneer factors, require continuous SWI/SNF activity to remain accessible and active. However, a subset of enhancers occupied by CTCF can employ alternate chromatin remodeler SNF2H to sustain accessibility, even during prolonged absence of SWI/SNF activity.

### A majority of gene promoters recover from loss of SWI/SNF activity

We then turned our attention to promoters, which have often been considered insensitive to SWI/SNF activity, or even to be repressed by SWI/SNF-mediated remodeling.^3, 12, 20^ Strikingly, analysis of promoter accessibility after 2 h of BRM014 treatment revealed a global reduction in chromatin accessibility, like that observed at enhancers (Figure 2A, 97% of promoters affected). As at enhancers, the promoters most affected by BRM014 inhibition are those with the highest levels of BRG1 ChIP-seq signal in control mESCs (Figure 2A), consistent with reduced accessibility reflecting a direct effect. Inhibitor treatment causes a marked increase in BRG1 binding at promoters, with the peak of BRG1 occupancy coinciding with the first (+1) well-positioned nucleosome (Figures 2A and 2B). We conclude that SWI/SNF broadly opens chromatin at promoters and that the immediate effects of inhibiting BRG1 are very similar at promoter and enhancer loci.

**Figure 2.**
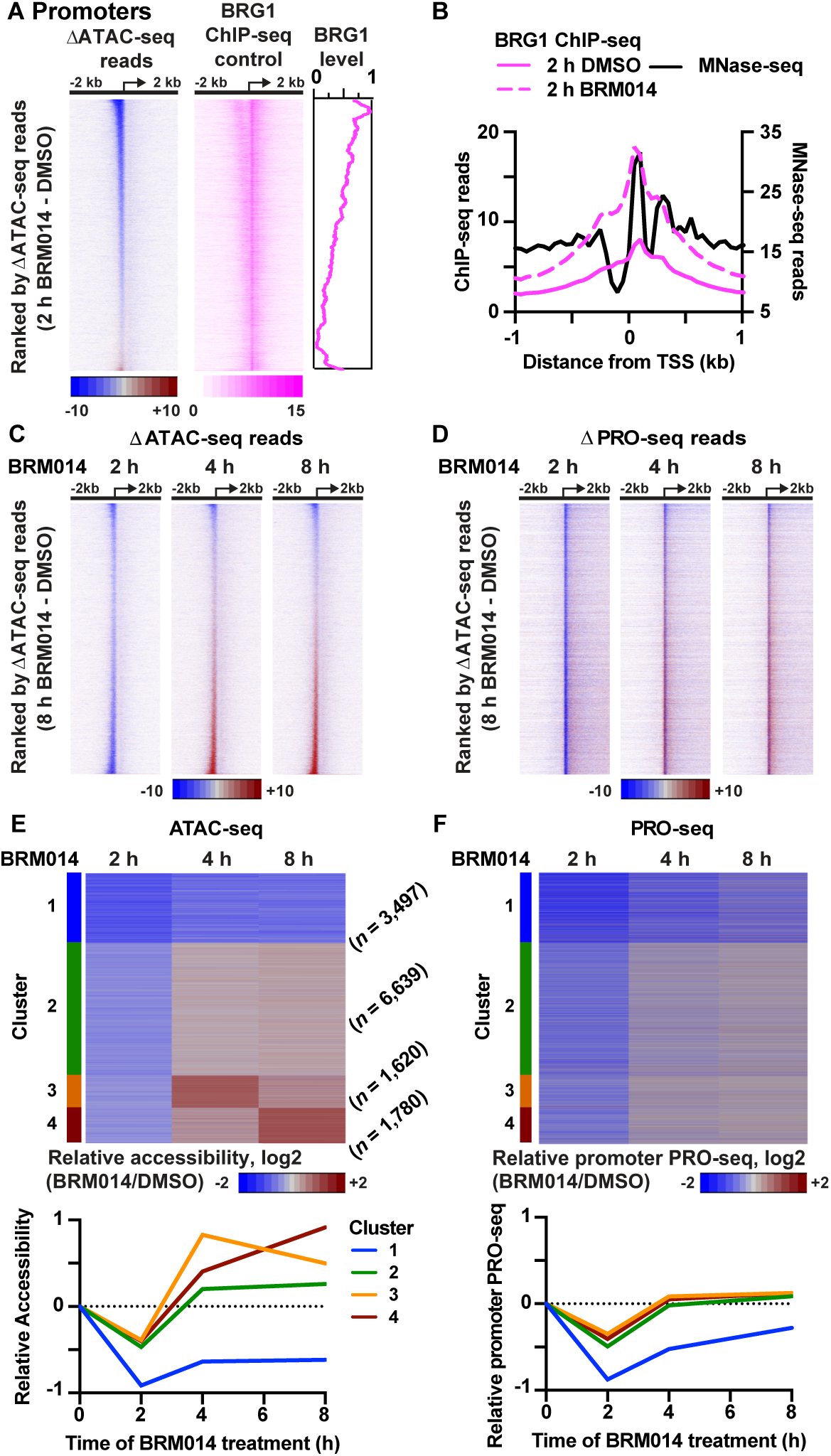
Promoters recover from SWI/SNF inhibition with variable kinetics. (**A**) Heatmaps showing the effects of 2 h BRM014 treatment on ATAC-seq signal at promoters (n = 13,536). Data are aligned to TSS. Sites are ranked by difference in ATAC-seq reads after 2 h BRM014 treatment (-450 to +149 bp from the TSS). BRG1 ChIP-seq signal is shown in the same rank order, as is relative BRG1 signal (summed from -750 to +249 bp relative to TSS). **(B)** Aggregate plot of quantitative BRG1 ChIP-seq signal around promoters in DMSO- and BRM014-treated cells. Average MNase-seq^28^ profile is shown to define the position of the NDR. Data are graphed in 50 bp bins. **(C and D)** Difference in ATAC-seq (C) and PRO-seq (D) signal after BRM014 treatment (compared to time-matched DMSO controls, as in A) for all promoters. Data were aligned to TSS and genes rank ordered by the difference in promoter ATAC-seq reads after 8 h treatment. **(E and F)** Clustering based on relative differences in ATAC-seq reads (as in C) defines four classes of responses to extended BRM014 treatment. The average value in each cluster for the relative ATAC-seq (-450 to +149 bp from the TSS) and PRO-seq (TSS to +150 nt) signals across the time course are shown at bottom See also Figure S3.

Extended BRM014 treatment, however, exhibited markedly different effects at promoters versus enhancers. In contrast to the persistent repression of accessibility observed at most enhancers, a majority of promoters effectively recover accessibility after 4 h of BRM014 treatment (Figure 2C). In fact, many promoters display even greater accessibility upon 4 h of SWI/SNF inhibition. These prominent accessibility changes were confirmed by ATAC-qPCR at selected loci (Figure S3A). The striking restoration of ATAC-seq signal at gene promoters following prolonged BRM014 treatment suggests that the loss of SWI/SNF activity can be functionally compensated at many promoters, and even over-compensated at some loci.

To determine how the observed changes in promoter chromatin accessibility impact gene transcription, we evaluated PRO-seq signals over the BRM014 treatment time course. After 2 h of BRM014 treatment, the widespread reduction of promoter accessibility was accompanied by a strong repression of transcription activity (Figure 2D), with a marked loss of promoter-proximal Pol II. These results are consistent with a requirement for accessible promoter chromatin to allow for transcription initiation. Upon longer SWI/SNF inhibition, as chromatin accessibility was restored at many gene promoters, transcription initiation and gene activity recovered concomitantly.

To investigate the variable promoter recovery during BRM014 treatment, promoters were clustered based on their chromatin accessibility changes over the BRM014 time course (Figure 2E). While most gene promoters (Clusters 2-4) were able to readily reinstate chromatin accessibility following the loss of SWI/SNF activity, about one-quarter of promoters (Cluster 1) remained repressed. Importantly, the inability of Cluster 1 genes to reinstate accessibility in the absence of BRG1 activity is reproducible and persistent, as these genes show substantially higher nucleosome occupancy in mESCs subjected to 24 h BRM014 treatment or following 72 h BRG1 knockout (Figure S3B and S3C).

Graphing promoter-proximal PRO-seq signal across the four clusters (Figure 2F) confirmed that changes in accessibility are generally mirrored by transcriptional changes. However, while promoters in Clusters 3 and 4 show evidence of elevated ATAC-seq signal at the 4 or 8 h time point as compared to DMSO controls, we find no evidence that transcription is broadly increased above control levels under these conditions (Figure 2F and below). Overall, these findings indicate that accessible promoter chromatin is necessary, but not sufficient, for gene transcription. Further, they support a model wherein the direct, immediate consequence of SWI/SNF inhibition is reduced accessibility and transcription at both promoters and enhancers. Whereas most promoters can compensate for loss of SWI/SNF activity to reestablish accessible chromatin and gene expression, a subset of promoters and a majority of enhancers are dependent upon SWI/SNF-mediated remodeling for appropriate accessibility and activity.

### Recovery from SWI/SNF inhibition is largely promoter-autonomous

We hypothesized that the variable ability of gene promoters to reinstate expression during prolonged BRM014 treatment might be connected to the activity of nearby enhancers. To test this model, we first stringently defined differentially expressed genes in 8 h BRM014-treated cells vs DMSO controls, using PRO-seq signal within gene bodies (TSS+250 to TES). This analysis revealed 633 downregulated genes and 324 upregulated genes (Fold-change > 1.5 and P adj < 0.001). As anticipated, Cluster 1 promoters were markedly enriched among genes with sustained downregulation of transcription as compared to all genes (Figure 3A). To confirm these gene sets, we assessed gene body PRO-seq levels over the BRM014 treatment time course (Figure 3B). For the downregulated genes, repression was notable at the earliest timepoint, suggesting that these genes are rapidly and persistently repressed by BRM014. In contrast, the upregulated genes showed gradually increased PRO-seq signal to a maximum at 8 h, suggesting that upregulation may occur more slowly following BRM014 treatment. For comparison, we defined a set of unchanged genes (FC < 1.1 and P adj > 0.5) that showed no appreciable differences in PRO-seq signal in BRM014-treated cells (Figure S4A).

**Figure 3.**
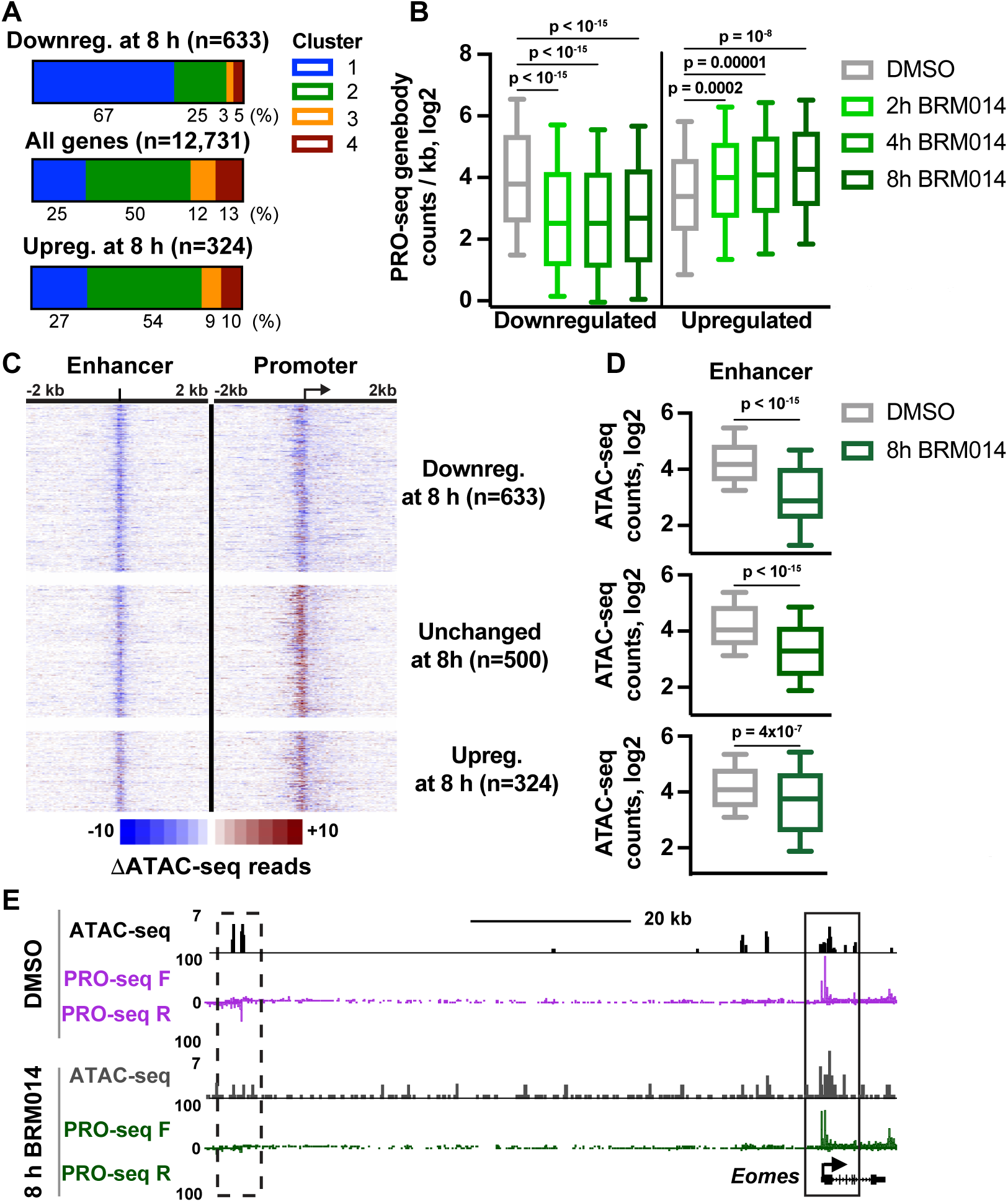
Recovery of gene expression during BRM014 treatment is not dependent on the activity of nearby enhancers. (**A**) Representation of promoter clusters among genes downregulated or upregulated after 8 h BRM014 treatment, as compared to all active genes longer than 1 kb. Percentages of genes in each cluster are indicated. **(B)** PRO-seq read density in gene bodies (TSS+250 to TES) is shown at downregulated and upregulated genes. Differentially expressed genes were those with a fold-change > 1.5 and an adjusted p-value < 0.001. Whiskers show the 10-90th percentiles and p-values are from Mann-Whitney test. **(C)** Heatmaps show the effects of 8 h BRM014 treatment on ATAC-seq signal at promoters (right) and their closest enhancers (left) for genes downregulated (n = 633), unchanged (subsampled, n = 500) and upregulated (n = 324) upon BRM014 treatment. Data are aligned to the enhancer center or gene TSS. **(D)** ATAC-seq counts at the closest enhancers (± 300 bp relative to the enhancer center) for the genes downregulated (top), unchanged (middle), or upregulated (bottom) after 8 h BRM014 treatment. Whiskers show the 10-90th percentiles and p-values are from Mann-Whitney test. **(E)** Genome browser image of ATAC-seq and PRO-seq data at the *Eomes* promoter (solid box) and associated enhancer (dashed box) in cells treated with BRM014 or DMSO for 8 h. See also Figure S4.

We then assessed chromatin accessibility at the nearest enhancer of the downregulated, unchanged, and upregulated gene sets, using heatmaps of ATAC-seq signal and box plot analyses of read counts (Figures 3C and 3D). At the downregulated genes (Figures 3C and 3D, top), consistent with their enrichment for Cluster 1 genes, promoter chromatin accessibility remained reduced after 8 h BRM014 treatment. Enhancers associated with downregulated genes also remained significantly repressed. For unchanged genes, where promoter chromatin accessibility was restored or even increased by 8 h BRM014, we observed persistent repression of the nearest enhancers (Figures 3C and 3D, middle). This result suggests that recovery of chromatin accessibility and gene activity at promoters can occur independently of enhancer inputs. Indeed, investigation of individual loci with well-defined enhancers that are essential for maintaining expression in mESCs, such as the *Eomes* gene^37^ (Figure 3E, validated enhancer shown in dashed box), demonstrates that gene activity is fully restored after 8 h of BRM014 treatment, despite continued reduction of both accessibility and eRNA synthesis at the cognate enhancer. Even at upregulated genes, which showed continually increasing activity during BRM014 treatment (Figure 3B), we find only partial recovery at nearby enhancers, with 38 % of the associated enhancers recovering to starting accessibility levels (Figures 3C and 3D, bottom). Thus, even genes that have increased expression following BRM014 treatment are generally near enhancers with lower accessibility and activity.

To address the relationship between enhancer and nearby promoter recovery in a different way, we divided enhancers into quartiles based on the level of accessibility after 8 h of BRM014 treatment and assessed ATAC-seq signal at the promoters nearest these enhancers. This analysis provided no evidence that enhancer recovery affects the reinstatement of accessibility at nearby promoters (Figure S4B). Indeed, promoters associated with the most persistently repressed enhancers were just as capable of restoring accessibility during extended BRM014 treatment as were promoters associated with enhancers that recovered entirely. We conclude that the restoration of accessibility and activity at gene promoters occurs autonomously of nearby enhancers. Notably, a similar, widespread disruption of enhancer-promoter communication following SWI/SNF perturbation was recently documented in prostate cancer cells.^25^ Together, these findings suggest that enhancer dysfunction is a general feature of prolonged SWI/SNF inhibition, that can occur in healthy as well as diseased cells.

### SWI/SNF-dependent promoters have chromatin features that are characteristic of enhancers

We then sought to define the features that discriminate SWI/SNF-dependent, Cluster 1 promoters from those that can compensate for loss of SWI/SNF activity. Investigation of chromatin architecture revealed that Cluster 1 promoters are characterized by lower average accessibility and exhibit particularly small and weak NDRs as compared to Cluster 2-4 promoters (Figures 4A and 4B). Analysis of PRO-seq data showed that Cluster 1 genes displayed lower occupancy by engaged Pol II in both sense and antisense directions (Figure 4C), as well as lower levels of RNA expression (Figure S5A). Cluster 1 genes are enriched for GO terms associated with cell signaling, development, and specific cell types or developmental lineages (Figure S5B). Notably, Cluster 1 is enriched for genes involved in neuron development and the cardiovascular system, lineages known to require BRG1,^38–40^ suggesting that BRG1 helps to poise these genes in mESCs for activation during development. Consistent with their enrichment in developmental genes, 25% of Cluster 1 genes were considered bivalent,^33^ as compared to 13% of all expressed genes (Figure S5C). However, 75% of Cluster 1 genes were not bivalent, indicating that bivalency and SWI/SNF dependence are distinct.

**Figure 4.**
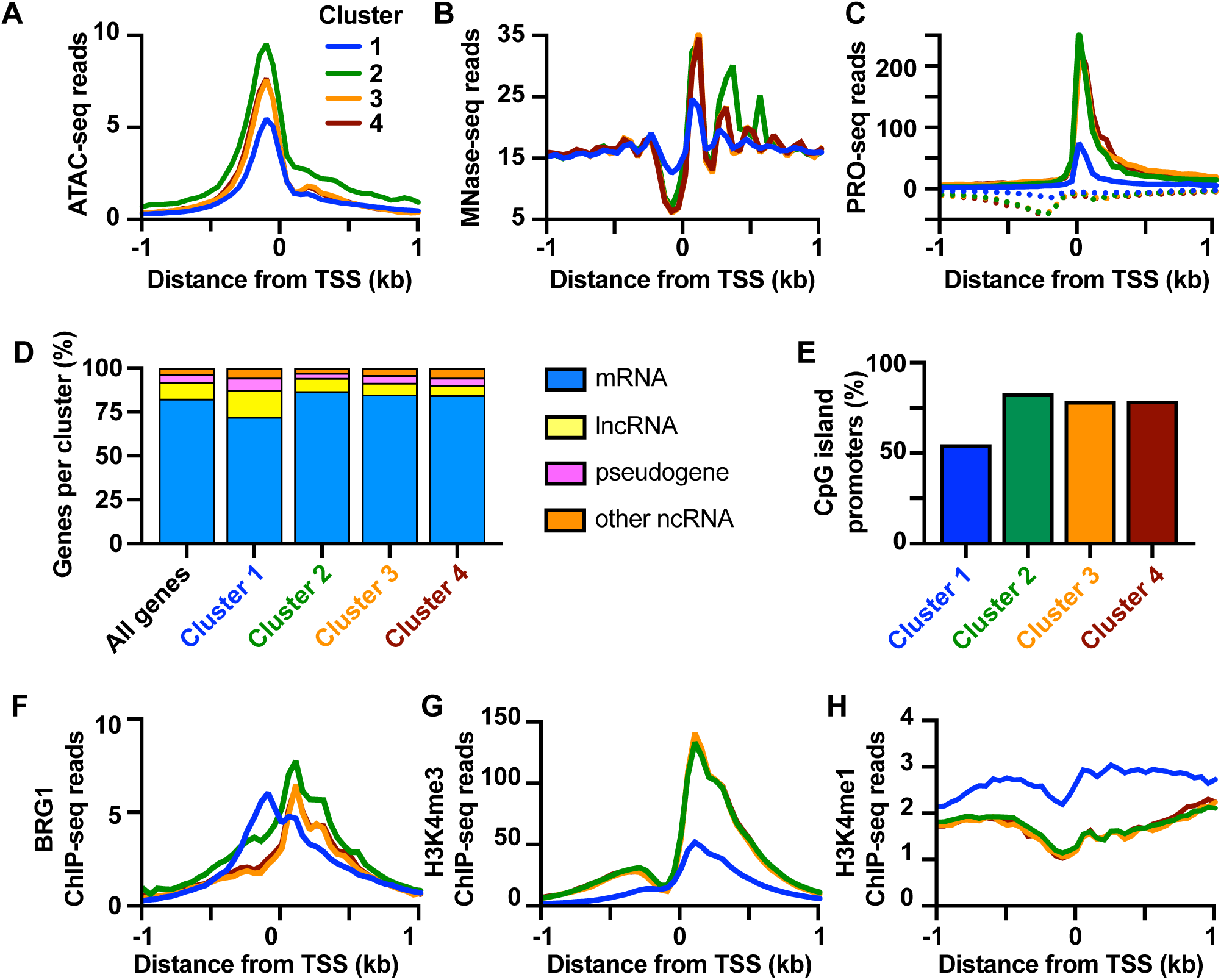
Promoters that are sensitive to SWI/SNF inhibition have distinct epigenetic characteristics. (A-C) Aggregate plots of average reads at promoters by cluster for ATAC-seq (A), MNase-seq^28^ (B) and PRO-seq (C) signal. Data are graphed in 50 bp bins. **(D)** Percentage of genes by transcript biotype for all annotated genes in each cluster. **(E)** Bar graph showing percentage of promoters by cluster overlapping a CpG island.^62^ **(F-H)** Aggregate plots of average reads at promoters by cluster for BRG1 ChIP-seq (F), H3K4me3 ChIP-seq^60^ (G) and H3K4me1 ChIP-seq^61^ (H) signal. Data are graphed in 50 bp bins. See also Figure S5.

Cluster 1 is enriched for non-coding RNA species, including lncRNAs, pseudogenes and pre-miRNAs (Figure 4D). Analysis of evolutionary conservation revealed that Cluster 1 promoters are less conserved on average than other promoters (Figure S5D). Cluster 1 promoters are less likely to overlap a CpG island than other clusters; however, more than 54% of Cluster 1 promoters are embedded in CpG islands (Figure 4E), and the profile of GC enrichment across Cluster 1 promoters is similar to that of Cluster 2-4 promoters (Figure S5E). These data thus demonstrate that SWI/SNF dependence is not dictated by GC content, as has been previously suggested,^41, 42^ and highlight that SWI/SNF inhibition can repress CpG-island promoters as well as those with lower GC content.

Strikingly, while BRG1 occupancy over most gene promoters is focused over the +1 nucleosome, Cluster 1 promoters are instead bound by BRG1 over the NDR (Figure 4F). This pattern is reminiscent of BRG1 localization at enhancers (Figure 1C), suggesting that BRG1 may be executing a shared, essential, activity at Cluster 1 promoters and enhancers. Characterization of histone modifications revealed that, in comparison to other genes, Cluster 1 promoters feature lower levels of H3K4me3 and significantly higher levels of H3K4me1 (Figure 4G, 4H). This finding is consistent with our determination that Cluster 1 genes tend to be lowly expressed, as levels of histone H3 modifications are known to reflect levels of transcriptional activity.^28, 43^ Moreover, this finding further emphasizes the similarities between non-recovering gene promoters and distal enhancers, which are generally characterized by the relative enrichment of H3K4me1 over H3K4me3. Overall, our data highlight the importance of SWI/SNF activity at genes with low expression, weak nucleosome depletion, and enhancer-like chromatin features.

### Promoter characteristics can predict gene expression changes following SWI/SNF perturbation in cancer cells

We next asked whether the distinguishing features of SWI/SNF-dependent promoters in mESCs, in particular the elevated H3K4me1 ChIP-seq and low ATAC-seq signals, could be used to predict gene responses to SWI/SNF inhibition in other systems (Figure 5A). We evaluated this in two types of cancer for which SWI/SNF is being pursued as a therapeutic target,^25, 26, 44, 45^ non-small cell lung cancer (NSCLC) and prostate cancer. Using existing H3K4me1 ChIP-seq and ATAC-seq data sets from A549 and H1299 NSCLC cells and LNCaP and VCaP prostate cancer cells (see STAR methods), we identified promoters within both the top 15 percent of H3K4me1 signal and bottom 15 percent of ATAC-seq signal. These promoters were predicted to be sensitive to SWI/SNF inhibition, and thus repressed by loss of SWI/SNF activity. Conversely, genes within the bottom 15 percent of H3K4me1 signal and top 15 percent of ATAC-seq signal were predicted to recover activity during SWI/SNF perturbation, and thus to be resistant to long term changes in gene expression.

**Figure 5.**
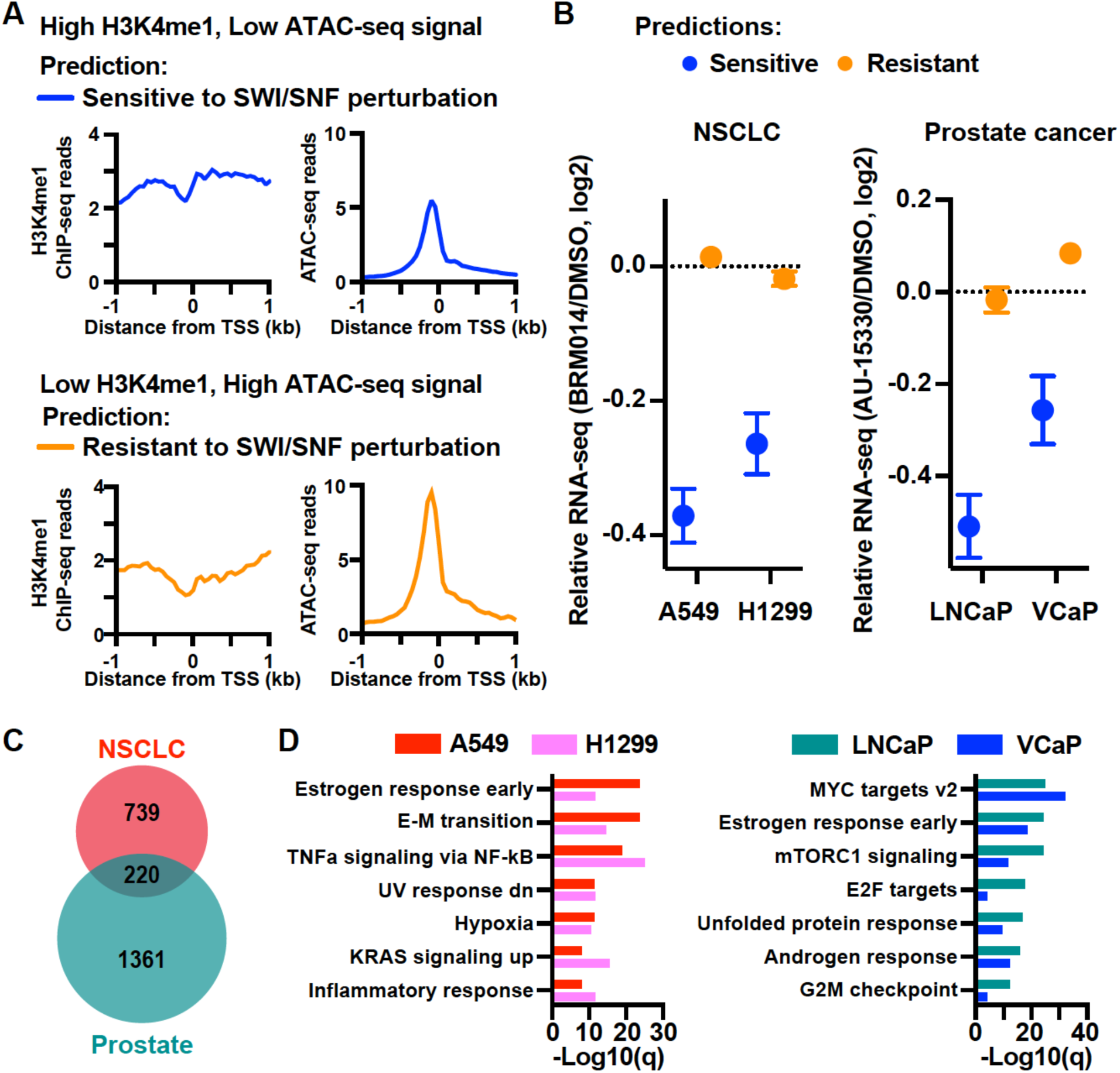
Epigenetic features of promoters can predict sensitivity to SWI/SNF inhibition or degradation. (**A**) Strategy to predict gene response to prolonged SWI/SNF disruption, using H3K4me1 ChIP-seq and ATAC-seq from control mESCs. Genes with high H3K4me1 ChIP-seq^61^ signal (top 15%) and low ATAC-seq signal (bottom 15%) were predicted to be sensitive while genes with low H3K4me1 ChIP-seq^61^ signal (bottom 15%) and high ATAC-seq signal (top 15%) are predicted to be resistant to SWI/SNF perturbation. H3K4me1 ChIP-seq and ATAC-seq signals were summed ± 500 bp relative to TSS. **(B)** Mean expression changes at genes predicted to be sensitive (blue) or resistant (orange) to SWI/SNF perturbation. The average log2 Fold-change in RNA-seq following SWI/SNF inhibition by 12 h BRM014 treatment (A549 and H1299 lung cancer cell lines, left) or 12 h degradation by AU-15330 (LNCaP and VCaP prostate cancer cell lines, right) is shown. Error bars represent SEM. See Methods for data sources and number of genes in each group. **(C)** Venn diagram showing the overlap between genes downregulated following BRM014 treatment in NSCLC cells (H1299 or A549) or following AU-15330 treatment in Prostate cancer cells^25^ (VCaP or LNCaP). Down regulated genes had a Fold-change > 1.5 and P adj < 0.001. P-value of the overlap calculated using the hypergeometric distribution was 7.3 e-22. **(D)** Top enriched Hallmark gene sets in genes downregulated (as in C) by BRM014 treatment in lung cancer lines (left) or AU-15330 treatment in prostate cancer lines^25^ (right). See also Figure S5.

We then performed RNA-seq on A549 and H1299 NSCLC cells treated with BRM014 for 12 h at 5 µM, a concentration at which cell growth was not affected (Figure S5F). In both NSCLC lines, genes predicted to be SWI/SNF-sensitive were indeed downregulated following SWI/SNF inhibition, whereas genes predicted to be resistant to BRM014 were unchanged (Figure 5B, left). We next analyzed published RNA-seq data from LNCaP and VCaP cells treated for 12 h with the BRG1/BRM PROTAC degrader AU-15330.^25^ Notably, the SWI/SNF ATPases are not simply inhibited in this system, but these proteins are instead rapidly degraded, providing an orthogonal method to test the generality of our predictions. Again, genes predicted to be sensitive to SWI/SNF activity based on their chromatin signatures showed significant downregulation upon loss of the SWI/SNF ATPases, whereas genes predicted to recover activity displayed unchanged activity following BRG1/BRM degradation (Figure 5B, right). Importantly, these data demonstrate that features associated with SWI/SNF-dependence in mESCs can accurately predict gene responses to SWI/SNF inhibition or degradation in markedly different cellular contexts. Furthermore, we could predict downregulated genes following SWI/SNF perturbation using ATAC-seq data alone, although with reduced accuracy (Figure S5G). Thus, using commonly available genomic data sets, we can successfully predict which genes will be most sensitive to SWI/SNF inhibition, a powerful possibility given the high-level interest in suppressing SWI/SNF activity in cancer.

That H3K4me1 and ATAC-seq signals, which vary across cell types, can predict gene sensitivity to SWI/SNF perturbation suggests that SWI/SNF-dependence is determined by the chromatin state at gene promoters, rather than being hard-wired by DNA sequence. Indeed, genes downregulated following SWI/SNF perturbation in NSCLC cells (A549 or H1299) differ substantially from those affected in prostate cancer lines (LNCaP or VCaP), despite significant overlap (Figure 5C, S5H-I). Moreover, gene ontology analysis of downregulated genes revealed largely different pathways repressed following long term SWI/SNF perturbation (Figure 5D).

Thus, in agreement with the variability in gene targets affected by SWI/SNF disruption in disease states, we find that genes repressed by sustained perturbation of SWI/SNF exhibit cell type specificity. Consequently, SWI/SNF dependence of gene expression cannot be predicted merely by sequence content. Our work reveals, however, that SWI/SNF dependence can be inferred by the chromatin state (ATAC-seq and H3K4me1) at gene promoters.

### EP400/TIP60 drives recovery of gene activity at most gene promoters

The above data suggest that Cluster 1 promoters lack a compensatory remodeler that enables recovery of chromatin accessibility following inhibition of SWI/SNF. To probe this possibility, we investigated ChIP-seq localization for several chromatin remodelers in mESCs. We found many remodelers to be present at similar levels across promoter clusters regardless of recovery capacity (Figure 6A), including SNF2H, which was implicated in the recovery of accessibility at CTCF-bound enhancers (Figure S2).

**Figure 6.**
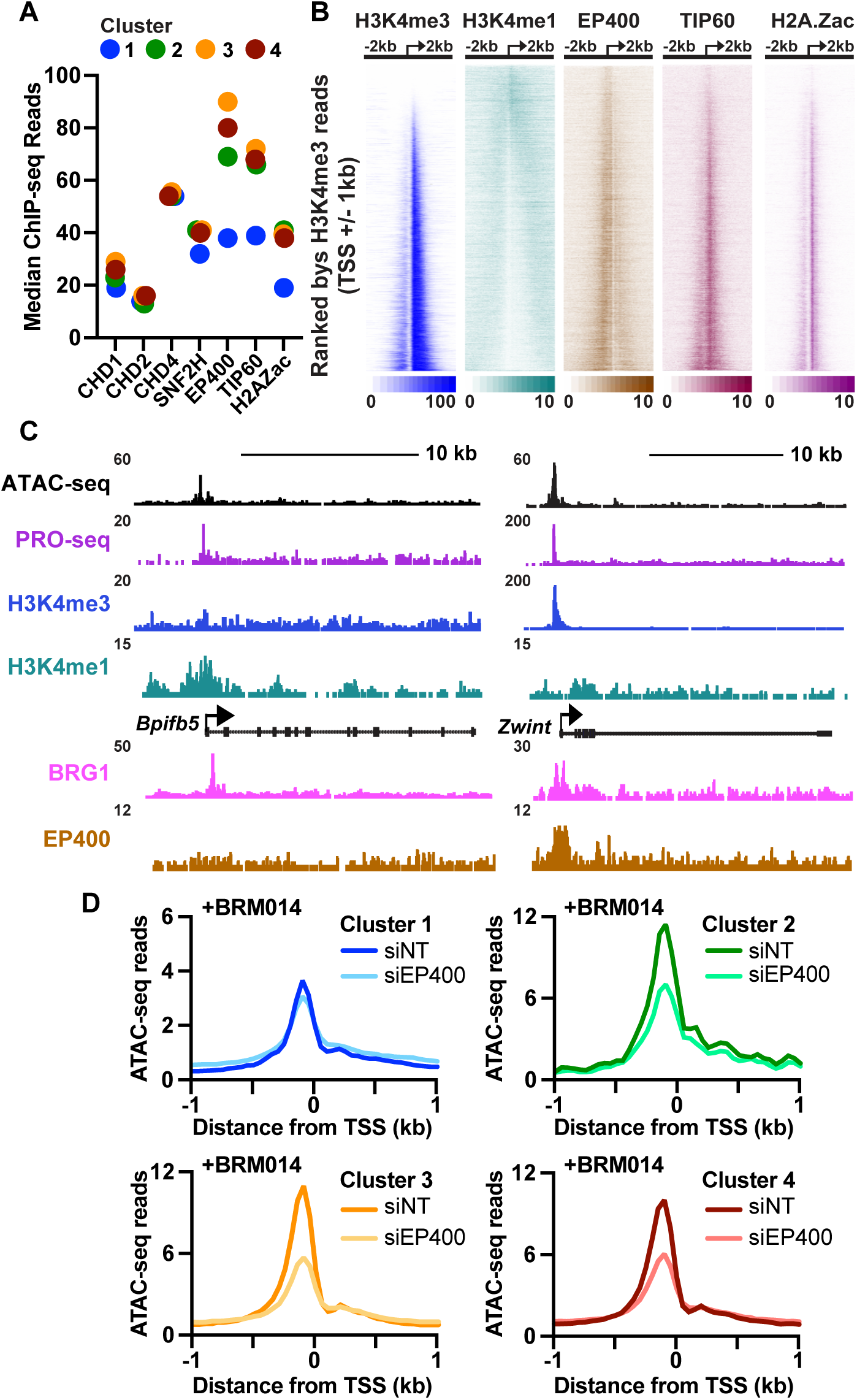
Recovery of accessibility at promoters following SWI/SNF inhibition is dependent on EP400/TIP60. (**A**) Median ChIP-seq signal (± 500 bp relative to TSS) for CHD1,^3^ CHD2,^3^ CHD4,^3^ SNF2H,^63^ EP400,^3^ TIP60^64^ and H2A.Zac^65^ across each promoter cluster. **(B)** Heatmaps of H3K4me3,^60^ H3K4me1,^61^ EP400,^3^ TIP60^64^ and H2A.Zac.^65^ Data are aligned to TSS and genes rank ordered by promoter H3K4me3 signal (± 1kb around TSS). **(C)** Genome browser images of representative SWI/SNF sensitive gene *Bpifb5* (left) and resistant gene *Zwint* (right). **(D)** Aggregate plots of ATAC-seq signal at promoters in each cluster, following 4 h BRM014 treatment under siNT conditions (n ≥ 2 per condition), graphed in 50 bp bins. See also Figure S6.

However, Cluster 1 genes were strongly depleted of both EP400 and TIP60, key subunits of the EP400/TIP60 complex (Figure 6A, S6A-B).^46^ The EP400/TIP60 complex both deposits and acetylates histone H2A.Z (H2A.Zac), such that H2A.Zac is a specific marker of complex activity in mammalian cells.^47^ Accordingly, we found that H2A.Zac was significantly depleted from Cluster 1 promoters (Figure 6A, S6A-B) as compared to promoters in Clusters 2-4, validating that EP400/TIP60 is preferentially localized to the promoters that recover from SWI/SNF inhibition. This finding was intriguing in light of previous reports that EP400/TIP60 binds H3K4me3 through its ING3 subunit.^48–50^ Selective recruitment of EP400/TIP60 to promoters enriched in H3K4me3 would thus provide a mechanistic explanation for the localization of this complex at Cluster 2-4 promoters. Indeed, heatmaps of all active promoters rank ordered by increasing H3K4me3 levels show a clear relationship between the H3K4me3 modification and levels of EP400, TIP60, and H2A.Zac (Figure 6B).

Based on our evaluation of individual genes (Figure 6C), as well as earlier reports that BRG1 and EP400 may work together to regulate chromatin accessibility at a set of promoters in mESCs,^3^ we tested whether EP400 enables efficient recovery of accessibility at Cluster 2-4 genes following BRM014 treatment. We performed siRNA knockdown of EP400 for 72 h, which was confirmed to substantially reduce both EP400 mRNA and protein levels (Figure S6C and S6D), followed by ATAC-seq with spike in for accurate quantitation. In the absence of SWI/SNF inhibitors, knockdown of EP400 was not associated with appreciable changes to promoter chromatin accessibility at genes in Clusters 1-4 (Figure S6E). Notably, EP400 was previously suggested to selectively regulate bivalent genes in mESCS.^3^ However, analysis of EP400 ChIP-seq revealed that EP400 binding is at background levels at bivalent genes (Figure S6F). Further, EP400 knockdown had minimal effects on chromatin accessibility at bivalent genes (Figure S6G), suggesting that other remodelers dominate the profile of chromatin accessibility during normal mESC growth, even at bivalent loci.

ATAC-seq data from cells depleted of EP400 and treated with BRM014 for 4 h showed clear effects of EP400 siRNA (Figure 6D), supporting an increased role for EP400 following loss of SWI/SNF activity. Importantly, EP400 depletion strongly reduced chromatin accessibility at Cluster 2-4 promoters versus a subtle effect at Cluster 1 (Figures 6D, S6H and S6I). These findings were confirmed by ATAC-qPCR analysis at selected genes (Figure S6J). Together, these data provide strong evidence that the activity of EP400/TIP60 enables the recovery of accessibility at Cluster 2-4 promoters in the absence of SWI/SNF activity. Mechanistically, this implies that the role of SWI/SNF at many promoters may be functionally compensated by EP400/TIP60. Cluster 1 promoters, which fail to effectively recruit EP400, would thus remain persistently repressed by BRM014 treatment.

### EP400/TIP60 loss creates a dependency on SWI/SNF in NSCLC

Our findings suggest that EP400/TIP60 may be critical for the establishment of appropriate chromatin architecture in cells lacking functional SWI/SNF. Indeed, analysis of TCGA data from NSCLC, for which mutations in SWI/SNF subunits are common, reveals that mutations in *EP400* are mutually exclusive with mutations in SWI/SNF subunits *BRG1* and *ARID1A* (Figure 7A). To directly test whether loss of *EP400* is synthetically lethal with disruption of SWI/SNF, we used CRISPR-Cas9 editing to introduce homozygous loss of function mutations into *EP400* in the NSCLC cell line A549 (Figures S7A and S7B). The *EP400-*KO cell line recapitulated a previously described increase in expression of epithelial-mesenchymal transition markers (Figure S7C), consistent with the enrichment of *EP400* mutations in metastatic tumors.^51^

**Figure 7.**
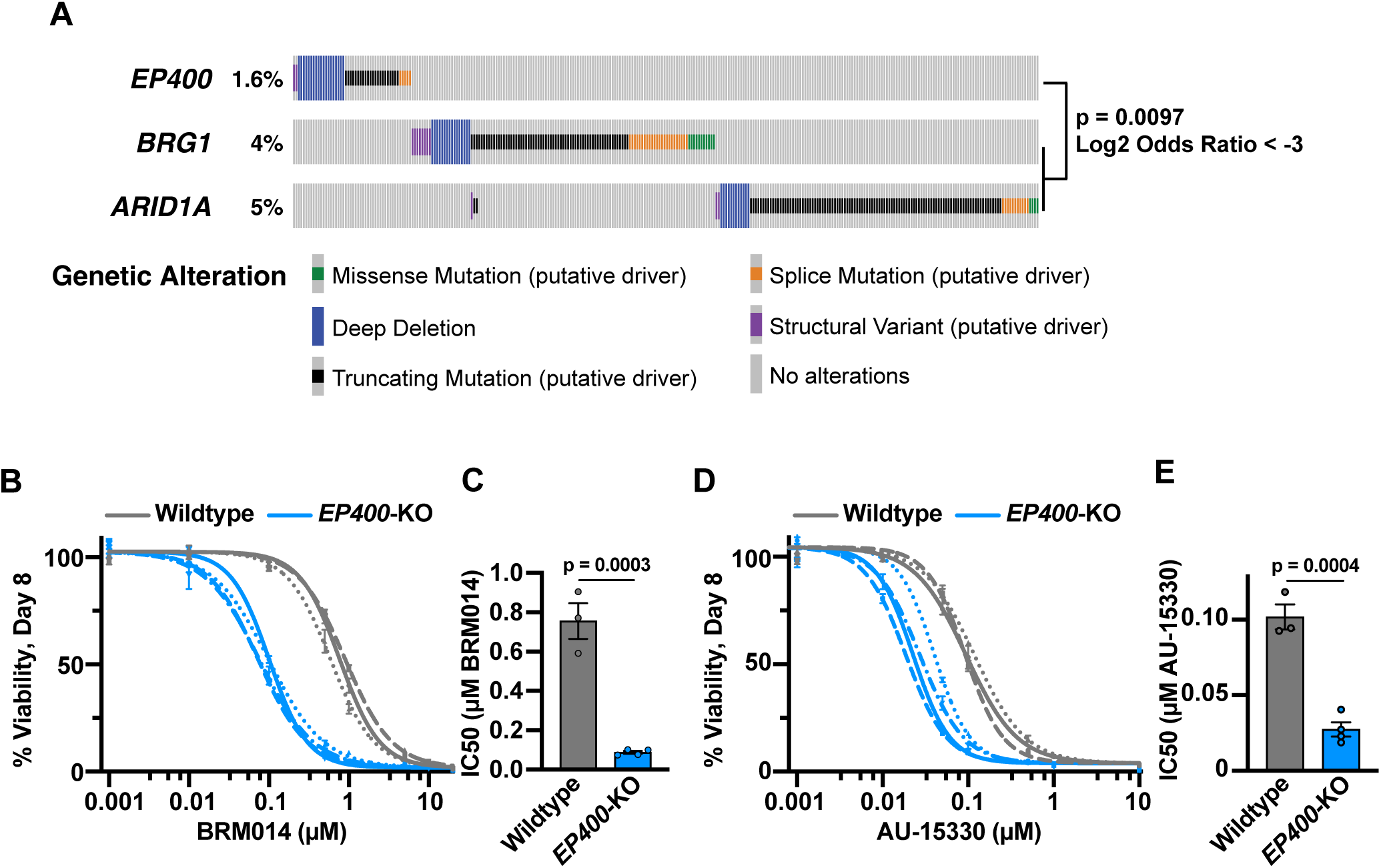
SWI/SNF activity is essential in non-small cell lung cancer cells lacking EP400. (**A**) NSCLC mutation data accessed through the cBio Portal (n = 3,311). Mutations of unknown significance were removed, and only samples profiling all 3 genes were analyzed. Fisher’s exact test performed for mutually exclusive relationship between *EP400* and *BRG1/ARID1A* mutations. The percentage of samples with the indicated mutations are indicated. **(B)** Drug dose response curves of wildtype and *EP400*-KO NSCLC (A549) cells following 8 days of treatment with BRM014. Each curve represents an independent experiment of the indicated cell line (n = 3 for wildtype and n = 4 for *EP400*-KO). Errors bars represent the SEM of three technical replicates. **(C)** Quantification of IC50 values from the dose response curves plotted in B. Error bars represent SEM. Individual values plotted as circles. P-values calculated by t-test. **(D)** Drug dose response curves of wildtype and *EP400*-KO A549 cells following 8 days of treatment with AU-15330. Each curve represents an independent experiment of the indicated cell line (n = 3 for wildtype and n = 4 for *EP400*-KO). Errors bars represent the SEM of three technical replicates. **(E)** Quantification of IC50 values from the dose response curves plotted in D. Error bars represent SEM. Individual values plotted as circles. P-values calculated by t-test. See also Figure S7.

We then tested whether EP400 loss affects cell growth in the presence of BRM014 (Figures 7B and 7C). We found that *EP400*-KO A549 cells displayed dramatically increased sensitivity to inhibition of SWI/SNF by BRM014, with IC50 values dropping from 756 nM to 87 nM. To perturb SWI/SNF function using an orthogonal approach, we tested whether the *EP400-* KO cells also displayed increased sensitivity to the PROTAC degrader of BRG1/BRM, AU-15330. Indeed, loss of EP400 sensitized cells to treatment with AU-15330, with IC50 values decreasing from 102 nM to 27 nM (Figures 7D and 7E). These results thus demonstrate that EP400 loss markedly sensitizes cells to perturbation of SWI/SNF activity through either a catalytic inhibitor or a PROTAC degrader, providing strong genetic support for a model where EP400/TIP60 becomes essential for recovery of chromatin architecture when SWI/SNF function is perturbed.

## DISCUSSION

Our data indicate that compensation by EP400/TIP60 masks a global role for SWI/SNF in promoting chromatin accessibility. We propose a model wherein SWI/SNF functions ubiquitously and continuously at nearly all promoters and enhancers to mobilize nucleosomes and enable binding of transcription factors and the general transcription machinery. In our mESCs, which lack mutations in SWI/SNF or other remodelers, this activity of SWI/SNF is sufficient to independently maintain open chromatin. Therefore, loss of EP400/TIP60 elicits little change in accessibility or gene activity under these conditions (Figure S6E,^3, 52, 53^). However, perturbation of SWI/SNF unveils a role for EP400/TIP60, revealing that EP400/TIP60 can reestablish accessibility at most gene promoters. This model provides a mechanistic explanation for the minor effects on gene activity observed upon disruption of SWI/SNF or EP400/TIP60 alone (Figure 3A,^3, 20, 21, 25^), and highlights the power of fast-acting inhibitors in assigning direct functions and untangling compensatory mechanisms. Critically, our work demonstrates that promoters and enhancers that are persistently repressed following perturbation of SWI/SNF do not represent the most direct targets of this remodeler; instead, they represent the sites at which compensation for SWI/SNF loss does not occur.

We find that the recovery of promoter accessibility and gene activity following SWI/SNF inhibition occurs largely independently of nearby enhancer activity. This finding is consistent with recent work in prostate cancer cells demonstrating that SWI/SNF degradation uncouples enhancer-promoter communication.^25^ Collectively, these results suggest that a common consequence of SWI/SNF perturbation could be promoter-autonomous gene activity. We suggest that the reduced input from enhancer loci on promoter activity could contribute to the altered gene expression profiles in cells with prolonged disruption of SWI/SNF activity.

The ability for cells to compensate for SWI/SNF loss is relevant in disease, where SWI/SNF is frequently mutated and is being explored as a therapeutic target. Our work reveals that the gene promoters most sensitive to loss of SWI/SNF activity have distinct features with prognostic value. Intriguingly, the promoters that fail to recover activity (Cluster 1) are those with weak accessibility and an enrichment of H3K4me1 over H3K4me3, features that are shared by enhancers. The absence of EP400/TIP60 at the persistently repressed promoters and enhancers supports that EP400/TIP60 is in part recruited by interactions with H3K4me3. The direct relevance of the molecular features we define at SWI/SNF dependent genes in mESCs is demonstrated by our ability to predict whether a gene will be sensitive or resistant to SWI/SNF perturbation in diverse cancer cell lines, using only ATAC-seq and H3K4me1 ChIP-seq data at gene promoters (Figure 5B).

Despite the ability of both SWI/SNF and EP400/TIP60 to increase promoter chromatin accessibility, these complexes possess distinct biochemical activities. SWI/SNF generates DNA accessibility through nucleosome sliding or eviction, whereas EP400/TIP60 exchanges H2A for H2A.Z and acetylates histone tails, including on H2A.Z. H2A.Z-containing nucleosomes have been reported to be hyper-labile,^54, 55^ reducing the nucleosomal barrier to transcription by Pol II, and acetylation of H2A.Z is tightly linked to transcription activation.^56–59^ Understanding how the disparate activities of SWI/SNF and EP400/TIP60 converge to enable promoter opening and transcription activation merits future investigation.

The synthetic lethality observed between EP400 and SWI/SNF in cancer patient data (for NSCLC) uncovers a dependency that could be targeted in cancer therapies, as the redundancy between SWI/SNF and EP400/TIP60 buffers the transcriptional response of cells against loss of either remodeler. Accordingly, our experiments demonstrate that EP400 mutations in lung cancer cells create a tumor-specific dependency on SWI/SNF, which may widen the therapeutic window for SWI/SNF-targeting compounds. Additionally, in pan-cancer analysis, EP400 mutations are enriched in metastatic tumors,^51^ and thus EP400 mutations may represent attractive indicators for targeting SWI/SNF more generally. For the SWI/SNF complex, mutations occur in over 20% of cancers and are frequently found to be driver mutations. Considering the prevalence of SWI/SNF mutations, we propose that inhibitors of the EP400/TIP60 complex present an attractive and unexplored therapeutic approach.

## Acknowledgments

We thank Fred Winston, Bob Kingston, Konrad Hochedlinger and Marie Bao for helpful discussions on this work. We also thank Zainab Jagani and the Novartis Institute for Biomedical Research for providing BRM014 under MTA and access to data from human lung cancer cells. This work was supported by the Van Maanen Fellowship from Harvard Medical School and a National Science Foundation Graduate Research Fellowship under Grant No. (DGE1745303) to EFA, a CIHR Banting Postdoctoral Fellowship BJEM and the Ludwig Center at Harvard (to KA and BJEM).

## Author contributions

Conceptualization: KA, EFA, BJEM. Methodology: KA, EFA, BJEM. Investigation: EFA, BJEM. Data analysis: EFA, BJEM, KA. Visualization: EFA, KA, BJEM. Project administration: KA. Funding acquisition: KA. Supervision: KA. Writing: EFA, BJEM, KA.

## STAR METHODS

## KEY RESOURCES TABLE

| REAGENT or RESOURCE | SOURCE | IDENTIFIER |
| --- | --- | --- |
| **Antibodies** |  |  |
| Rabbit monoclonal, BRG1 | Cell Signaling Technology | Cat # 49360, RRID:AB_2728743 |
| Rabbit monoclonal, BAF155 | Cell Signaling Technology | Cat # 11956, RRID:AB_2797776 |
| Rabbit monoclonal, ARID1A | Cell Signaling Technology | Cat # 12354, RRID:AB_2637010 |
| Rabbit polyclonal, EP400 | Abcam | Cat # ab70301, RRID:AB_1269644 |
| Rabbit polyclonal, H3 | Abcam | Cat # ab1791, RRID:AB_302613 |
| Rabbit monoclonal, Brg1 | Abcam | Cat # ab10641 |
| **Chemicals, peptides, and recombinant proteins** |  |  |
| BRM014 | Novartis Institutes for BioMedical Research | n/a |
| LIF | Cell Guidance Systems | Cat # GFM200 |
| PD0325901 | Reprocell, | Cat # 04-0006 |
| CHIR99021 | Reprocell, | Cat # 04-0004 |
| Biotin-11-NTPs | Perkin Elmer | Cat # NEL54(2/3/4/5)001 |
| AU-15330 | MedChem Express | Cat # HY-145388 |
| Protease Inhibitor Cocktail | Roche | Cat # 11836170001 |
| Lipofectamine RNAiMAX | Thermo Fisher | Cat # 13778075 |
| TRIzol reagent | Invitrogen | Cat # 15596018 |
| Trypan Blue | VWR | Cat # AAA18600-14 |
| DSG crosslinker | Cova Chem | Cat # 13301-5x100 |
| Formaldehyde | Sigma-Aldrich | Cat # F8775 |
| Protein A agarose beads | Millipore | Cat # 16-125 |
| Proteinase K | NEB | Cat # P8107 |
| Phenol:chloroform:isoamyl alcohol | Sigma-Aldrich | Cat # P3803 |
| IGEPAL CA-630 | Sigma-Aldrich | Cat # I8896 |
| SUPERase-In | Thermo Fisher | Cat # AM26976 |
| Dynabeads MyOne Streptavidin C1 | Thermo Fisher | Cat # 65001 |
| T4 RNA Ligase I | NEB | Cat # M0437 |
| RNA 5' Pyrophosphohydrolase | NEB | Cat # M0356 |
| T4 PNK Reaction Buffer | NEB | Cat # B0201 |
| T4 PNK | NEB | Cat # M0201 |
| SuperScript IV Reverse Transcriptase | Thermo Fisher | Cat # 18090010 |
| ProNex Size-Selective Purification System | Promega | Cat # NG2001 |
| DNase I, amplification grade | Invitrogen | Cat # 18068015 |
| **Critical commercial assays** |  |  |
| NEBNext Ultra II DNA Library Prep Kit for Illumina | NEB | Cat # E7103S |
| Illumina TruSeq Stranded Total RNA Library Prep Gold | Illumina | Cat # 20020598 |
| MinElute PCR Purification Kit | Qiagen | Cat # 28004 |
| NEBNext High-Fidelity 2X PCR Master Mix | NEB | Cat # M01541 |
| Total RNA Purification Kit | Norgen Biotek | Cat # 17250 |
| RNeasy Mini Kit | Qiagen | Cat # 74104 |
| Direct-zol RNA miniprep kit | VWR | Cat # 76211-340 |
| CellTiter-Glo 2.0 Assay | Promega | Cat # G9242 |
| **Deposited data** |  |  |
| Raw and analyzed data | This paper | GEO: GSEXXXXX |
| mESC MNase-seq | Henriques et al.<sup>28</sup> | GEO: GSE85191 |
| mESC TT-seq | Vlaming et al.<sup>60</sup> | GEO: GSE178230 |
| MNase-seq from control and BRM014 treated mESCs | Iurlaro et al.<sup>21</sup> | GEO: GSE158345 |
| ATAC-seq from control and *BRG1*-KO mESCs | King and Klose<sup>8</sup> | GEO: GSE87822 |
| ATAC-seq from control and *SNF2H*-KO mESCs | Barisic et al.<sup>4</sup> | GEO: GSE112130 |
| mESC H3K27ac ChIP-seq | Vlaming et al.<sup>60</sup> | 4DN:<br>4DNESQ33L4G7 |
| mESC H3K4me3 ChIP-seq | Vlaming et al.<sup>60</sup> | GEO: GSE178230 |
| mESC H3K4me1 ChIP-seq | Buecker et al.<sup>61</sup> | GEO: GSE56098 |
| mESC H2A.Z and H2A.Zac ChIP-seq | Hu et al.<sup>65</sup> | GEO: GSE34483 |
| mESC CHD1, CHD2, CHD4, and EP400 ChIP-seq | Dieuleveult et al.<sup>3</sup> | GEO: GSE64825 |
| mESC TIP60 ChIP-seq | Ravens et al.<sup>64</sup> | GEO: GSE69671 |
| mESC CTCF ChIP-seq | Justice et al.<sup>66</sup> | GEO: GSE137272 |
| mESC SNF2H ChIP-seq | Song et al.<sup>63</sup> | GEO: GSE123670 |
| mESC OCT4, SOX2, and NANOG ChIP-seq | King and Klose<sup>8</sup> | GEO: GSE87822 |
| LNCaP and VCaP RNA-seq and ATAC-seq | Xiao et al.<sup>25</sup> | GEO: GSE171523 |
| VCaP H3K4me1 ChIP-seq | Baumgart et al.<sup>67</sup> | GEO: GSE148400 |
| LNCaP H3K4me1 ChIP-seq | Sugiura et al.<sup>68</sup> | GEO: GSE122922 |
| A549 H3K4me1 ChIP-seq | Dunham et al.<sup>69</sup> | GEO: GSE29611 |
| A549 ATAC-seq | Dunham et al.<sup>69</sup> | GEO: GSE169955 |
| H1299 H3K4me1 ChIP-seq | Suzuki et al.<sup>70</sup> | SRA: DRR016953 |
| H1299 ATAC-seq | Kim et al.<sup>71</sup> | GEO: GSE141060 |
| **Experimental models: Cell lines** |  |  |
| F121-9 | Jaenisch/Gribnau labs | 4DNSRMG5APUM |
| A549 cell line | ATCC | CCL-185 |
| H1299 cell line | ATCC | CRL-5803 |
| A549 *EP400*-KO | This paper |  |
| **Oligonucleotides** |  |  |
| Table S1 | This paper | N/A |
| siEP400, Dharmacon J-058750-12 | Dharmacon | J-058750-12 |
| Non-Targeting siRNA | Dharmacon | D-001210-02-05 |
| ERCC RNA Spike-In Mix | Thermo Fisher | Cat # 4456740 |
| **Software and algorithms** |  |  |
| bowtie 1.2.2 | Langmead et al.<sup>72</sup> |  |
| STAR 2.7.3a | Dobin et al.<sup>73</sup> |  |
| R 3.6.1 | www.r-project.org |  |
| Rsubread 2.0.1 | Liao et al.<sup>74</sup> |  |
| DESeq2 1.26.0 | Love et al.<sup>75</sup> |  |
| Prism | GraphPad |  |
| Partek Genomics Suite 6.16.0812 | www.partek.com |  |
| get_gene_annotations.sh | Martin et al.<sup>76</sup> |  |
| make_heatmap | Martin et al.<sup>77</sup> |  |
| trim_and_filter_PE.pl | Martin et al.<sup>77</sup> |  |
| bowtie2stdBedGraph.pl | Martin et al.<sup>77</sup> |  |
| cutadapt | Martin et al.<sup>78</sup> |  |
| dREG | Wang et al.<sup>79</sup> |  |
| Samtools 1.9 | Li et al.<sup>80</sup> |  |
| Kallisto 0.45.1 | Bray et al.<sup>81</sup> |  |
| HOMER 4.10.3 | Heinz et al.<sup>82</sup> |  |
| The Database for Annotation, Visualization and Integrated Discovery (DAVID) v6.8 | Huang et al.<sup>83,84</sup> |  |

## RESOURCE AVAILABILITY

### Lead contact

Further information and requests for resources and reagents should be directed to and will be fulfilled by the lead contact, Karen Adelman.

### Materials availability

Unique and stable reagents generated in this study are available upon request.

### Data and code availability

All PRO-seq, ChIP-seq, ATAC-seq, and RNA-seq data have been deposited at GEO and are publicly available as of the date of publication. Accession numbers are listed in the key resources table.

## EXPERIMENTAL MODEL AND SUBJECT DETAILS

### Cell Culture and Inhibitor Treatments

#### Cell Culture

F121-9-CASTx129 female mouse hybrid embryonic stem cells (mESCs) were obtained from David Gilbert (Florida State University) and cultured at 37°C in 5% CO_2_. Cells were maintained in serum-free ES medium (SFES) composed of 50% Neurobasal Media (Gibco 21103-049), 50% DMEM/F12 (Gibco 11320-033), 0.5X N2 Supplement (Gibco 17502-048), 0.5X B27(+RA) (Gibco 17504-044) and 0.05% BSA (Gibco 15260-037) and supplemented with 2 mM glutamine (Gibco 25030-081), 1.5x10^-4^ M monothioglycerol (Sigma M6145), 1 µM MEK inhibitor (PD03259010; Reprocell 04-0006-02), 3 µM GSK3 inhibitor (CHIR99021;Reprocell 04-0004-02), and 1,000 U/mL leukemia inhibiting factor (LIF; Cell Guidance Systems GFM200). *Drosophila* S2 cells were grown at 27°C in Shields and Sang M3 Insect Medium (Sigma S3652) supplemented with bactopeptone (Difco 2116), yeast extract (Sigma Y-1000), and 10% FBS (Invitrogen 16000044).

*A549* cells were obtained from the ATCC and cultured at 37C in 5% CO2. Cells were maintained in F12-K medium composed of F12-K +L-glutamine (Gibco 21127-022), 10% fetal bovine serum (Fisher Scientific 16-000-044) and Penicillin-Streptomycin (Thermo Fisher 15140163). *H1299* cells were obtained from the ATCC and cultured at 37C in 5% CO2. Cells were maintained in RPMI medium consisting of RPMI 1640 with L-glutamine (Corning MT 10-040-CV), 10% fetal bovine serum (Fisher Scientific 16-000-044) and Penicillin-Streptomycin (Thermo Fisher 15140163). All cells were tested routinely for mycoplasma contamination.

#### BRM014 Treatment

SWI/SNF inhibitor BRM014^19^ was provided by Novartis Institutes for BioMedical Research (Cambridge, MA) and was resuspended in dimethyl sulfoxide (DMSO) for a 10 mM stock. mESCs were treated at a final concentration of 1 µM. For viability experiments, a TC20 Automated Cell Counter (Bio-Rad) was used to collect cell counts in duplicate for both unstained cells and cells stained with Trypan Blue (VWR AAA18600-14). Average cell counts for each condition were used to generate cell growth and viability curves.

## METHOD DETAILS

### Western Blots

Whole cell extracts were resolved using a Novex™ WedgeWell™ 6% Tris-Glycine Mini Protein Gel (Thermo Fisher XP00065BOX). Samples were transferred to a polyvinylidene difluoride (PVDF) membrane (Bio-Rad 1620177). After blocking in 5% BSA, membranes were incubated overnight with primary antibodies: BRG1 (Cell Signaling Technology #49360), BAF155 (Cell Signaling Technology #11956), ARID1A (Cell Signaling Technology #12354), EP400 (Abcam #ab70301), or H3 (Abcam #ab1791). Membranes were incubated with horseradish-peroxidase-conjugated secondary antibodies (Jackson ImmunoResearch 111-035-144) before being visualized using SuperSignal™ West Pico PLUS Chemiluminescent Substrate (Thermo Fisher 34579) and the ChemiDoc Imaging System (Bio-Rad).

### ATAC-seq Library Preparation

#### Cell Preparation and Transposition

ATAC-seq was performed as described in,^85^ with some modifications. In brief, 1 x 10^5^ cells per sample were washed with ice-cold 1X PBS and centrifuged at 500 × *g* for 5 min at 4°C. Cells were then resuspended in 50 µL CSK Lysis Buffer (10 mM PIPES, pH 6.8, 100 mM NaCl, 300 mM sucrose, 3 mM MgCl_2_, 0.1% Triton-X-100), incubated on ice for 5 min, and centrifuged for 5 min at 500 × *g* and 4°C. To allow for downstream spike normalization, *Drosophila* S2 cells were harvested in parallel and processed as described above, with spin speeds increased to 1000 x *g*. For each reaction, 1 x 10^5^ mESCs and 5 x 10^4^ S2 cells were resuspended in Tagment DNA Buffer and treated with 3 µL TDE1 Tagment DNA Enzyme (Illumina 20034197). After thorough mixing, samples were incubated at 37°C for 30 minutes. Tagmented DNA was subsequently purified using a MinElute PCR Purification Kit (Qiagen 28004).

#### Library Preparation

Purified samples were combined with NEBNext High-Fidelity 2X PCR Master Mix (New England Biolabs M01541) for amplification. As described in,^85^ custom primers were used to incorporate Illumina adaptors and index sequences into sample fragments. Libraries were sequenced at The Bauer Core Facility at Harvard University on an Illumina NovaSeq using an S1 flow cell and a paired-end 100-bp cycle run.

#### ATAC-qPCR

To validate ATAC-seq results, qPCR was performed using experimental primers (Table S1) targeting a panel of candidate genes and enhancer regions, as well as a set of three ‘background’ primer pairs targeting nongenic regions of closed chromatin. For each sample, Cq values of experimental primers were normalized to the average Cq value across all background primers for that same sample, allowing differences in accessibility between conditions to be expressed in terms of “normalized accessibility.”

### ChIP-seq Library Preparation

#### Chromatin Isolation and Sonication

After the indicated treatment interval, cells were fixed for 1 h in 2 mM DSG with the addition of 1% formaldehyde for the final 12.5 min, as described in.^86^ Crosslinking was quenched by the addition of glycine to a final concentration of 0.125 M. Cells were collected and washed with ice-cold 1X PBS before being resuspended in Sonication Buffer (20 mM Tris, pH 8.0, 2 mM EDTA, 0.5 mM EGTA, 1X Complete Mini EDTA-free Protease Inhibitor Cocktail [Roche 11836170001], 0.5% SDS, and 0.5 mM PMSF) at a concentration of 1 x 10^8^ cells per mL. Chromatin was sheared to an average fragment size of ∼200 bp using a QSonica sonicator, flash-frozen in liquid nitrogen, and stored at -80°C until use.

#### Immunoprecipitation

ChIP material was diluted into IP buffer (20 mM Tris pH 8.0, 2 mM EDTA, 0.5% Triton X-100, 150 mM NaCl, 10% Glycerol, 5% BSA) and pre-cleared with 30 μL of Protein A agarose beads (Millipore Cat No. 16-125) for 1 h at 4°C. Cleared samples were collected and combined with 30 μL of primary antibody (Brg1: Abcam ab10641 - EPNCIR111A) before overnight incubation at 4°C with rotation. Subsequently, 200 μL Protein A beads were added to each IP reaction, and samples were rotated for 2 h at 4°C. Samples were washed once with Low-Salt Buffer (20 mM Tris pH 8.0, 2 mM EDTA, 1% Triton X-100, 150 mM NaCl, 0.1% SDS), three times with High-Salt Buffer (20 mM Tris pH 8.0, 2 mM EDTA, 1% Triton X-100, 500 mM NaCl, 0.1% SDS), once with Lithium Chloride Buffer (20 mM Tris pH 8.0, 2 mM EDTA, 250 mM LiCl, 1% IGEPAL CA-630, 1% sodium deoxycholate), and twice with TE Buffer (10 mM Tris-HCl, 1mM EDTA). Each wash was performed by rotating samples for 3 minutes with 1 mL volume of ice-cold wash solution. Two elutions were performed by resuspending beads in Elution Buffer (100 mM NaHCO_3_, 1% SDS) and rotating for 15 min at room temperature (22°C). The combined eluate was supplemented with 200 mM NaCl and incubated overnight in a 65°C water bath. Samples were treated with Proteinase K (New England Biolabs P8107), extracted with phenol:chloroform:isoamyl alcohol (25:24:1, Sigma P3803), and resuspended in 65 μl H_2_O. To enable accurate sample normalization, an equal amount of fragmented *D. melanogaster* DNA was added to the eluate of each sample.

#### Library Preparation

Libraries were prepared using the NEBNext Ultra II DNA Library Prep Kit for Illumina (New England Biolabs) according to the manufacturer’s instructions. Libraries were sequenced on an Illumina NovaSeq using an S1 flow cell and a paired-end 100-bp cycle run, with sequencing performed by The Bauer Core Facility at Harvard University.

### PRO-seq Library Preparation

#### Cell Permeabilization

Precision run-on sequencing (PRO-seq) was performed based on the protocol described in,^27^ with some modifications. All steps of PRO-seq sample preparation were performed on ice, and all buffers were thoroughly chilled on ice before being added to the reaction. Cells were released using Accutase, collected with ice-cold media, washed with PBS, and resuspended in 0.25 mL Buffer W (10 mM Tris-HCL pH 8.0, 10 mM KCl, 250 mM sucrose, 5 mM MgCl_2_, 1 mM EGTA, 0.5 mM DTT, 10% glycerol). Then 10 mL Buffer P (Buffer W + 0.1% IGEPAL CA-630 (Sigma I8896) was carefully added to each sample. Samples were incubated on ice for 5 min, then centrifuged at 4°C and 400 x *g* for 4 min. Permeabilized cells were resuspended in 10 mL of Buffer W and centrifuged at 4°C and 400 x *g* for 4 min, before being resuspended in Buffer F (50 mM Tris-CL pH 8.0, 5 mM MgCl_2_, 1.1 mM EDTA, 0.5 mM DTT, 40% glycerol, 1 µL/mL SUPERase-In (Thermo Fisher AM26976) at a final volume of 1 x 10^6^ permeabilized cells per 50 µL. Immediately after processing, samples were flash-frozen using liquid nitrogen and stored at -80°C.

#### Biotin Run-On and RNA Purification

For each sample, 1 x 10^6^ permeabilized mES cells were spiked with previously prepared permeabilized *Drosophila* S2 cells at a proportion of 5% to enable downstream data normalization. Permeabilized cells were then combined with 2X Run-On Master Mix (10 mM Tris-Cl pH 8.0, 300 mM KCl, 1% Sarkosyl, 5 mM MgCl_2_, 1 mM DTT, 200 µM biotin-11-A/C/G/UTP (Perkin-Elmer NEL544001EA / NEL542001EA / NEL541001EA / NEL543001EA), 0.8 U/µL SUPERase-In (Thermo Fisher AM26976) and incubated at 30°C for 5 min to allow the biotin-NTP run-on reaction to proceed. Following run-on, RNA was isolated using the Total RNA Purification Kit (Norgen Biotek 17250) according to the manufacturer’s instructions.

#### Library Preparation

Purified RNA was subject to chemical fragmentation with 2X RNA Fragmentation Buffer (150 mM Tris-Cl pH 8.3, 225 mM KCl, 9 mM MgCl_2_) for 5 min at 94°C. Chilled fragmented RNA was combined with 48 µL Dynabeads MyOne Streptavidin C1 (Thermo Fisher 65001) in Binding Buffer (300 mM NaCl, 10 mM Tris-HCl pH 7.4, 0.1% Triton-X-100) and rotated for 20 min at room-temperature. RNA-bound beads were washed two times each with High-Salt Buffer (2 M NaCl, 50 mM Tris-HCl pH 7.4, 0.5% Triton-X-100), Binding Buffer (described above), and Low-Salt Buffer (5 mM Tris-HCl pH 7.4, 0.1% Triton-X-100), then resuspended in TRIzol Reagent (Invitrogen 15596026). RNA was eluted from the beads via two sequential rounds of incubation, each for 5 min at 65°C. Chloroform extraction was used to purify isolated RNA. Purified RNA was resuspended in 10 µM VRA3 adaptor (/5Phos/rGrArUrCrGrUrCrGrGrArCrUrGrUrArGrArArCrUrCrUrGrArArC/3InvdT/) and treated with T4 RNA Ligase I (New England Biolabs M0437) for 2 h at room temperature (22°C) to enable 3’ adaptor ligation. Desired RNA species were captured with Dynabeads MyOne Streptavidin C1 in the presence of a blocking oligo (TCCGACGATCCCACGTTCCCGTGG/3InvdT), after which the beads were sequentially washed with High-Salt, Binding, Low-Salt, and 1X Thermo Pol (New England Biolabs B9004) Buffers. Beads were next resuspended in 1X Thermo Pol Buffer and treated with 2 µL RNA 5’ Pyrophosphohydrolase (New England Biolabs M0356) at 37°C for 1 h to promote decapping of 5’ RNA ends. Beads were washed in High-Salt Buffer and Low-Salt Buffer, then resuspended in 1X T4 PNK Reaction Buffer (New England Biolabs B0201). Samples were incubated at 37°C for 1 h after the addition of T4 PNK (New England Biolabs M0201) to allow 5’-hydroxyl repair. A second ligation step was performed as described above to ligate the VRA5 5’ RNA adaptor (rCrCrUrUrGrGrCrArCrCrCrGrArGrArArUrUrCrCrA). Beads were washed twice each with High-Salt, Binding, and Low-Salt Buffers, then washed once in 0.25X FS Buffer (12.5 mM Tris-HCl pH 8.3, 18.75 mM KCl, 0.75 mM MgCl_2_). Twenty-five pmol of RP1 primer (AATGATACGGCGACCACCGAGATCTACACGTTCAGAGTTCTACAGTCCGA) was added to samples, after which reverse transcription was performed using SuperScript IV Reverse Transcriptase (Thermo Fisher 18090010). Final library products were eluted by heating samples twice to 95°C for 30 sec each, then amplified by 12 cycles of PCR with primer RP1, Illumina TruSeq PCR primer RPI-X, and Phusion Polymerase (New England Biolabs M0530). The ProNex Size-Selective Purification System (Promega NG2001) was used at a 2.8X ratio to purify amplified libraries. Libraries were sequenced at The Bauer Core Facility at Harvard University on an Illumina NovaSeq using an S4 flow cell and a paired-end 100-bp cycle run.

### siRNA Transfection

#### Cell Culture

For EP400 knockdown experiments, mESCs were transfected with either a non-targeting control siRNA (siNT) or a commercially available on-target siRNA against mouse *Ep400* (Dharmacon J-058750-12) (siEP400) using Lipofectamine RNAiMAX Transfection Reagent (Thermo Fisher 13778075). Cells were maintained for 72 h before harvest.

#### Knockdown Validation

To ensure that effective knockdown of *Ep400* was achieved under the conditions described above, cells transfected with either non-targeting (siNT) or on-target (siEP400) siRNA were harvested after 72 h for analysis of mRNA and protein levels. To analyze mRNA expression, RNA was extracted using the RNeasy Mini Kit (Qiagen 74104). cDNA synthesis was performed using hexamer primers and SuperScript IV Reverse Transcriptase (Thermo Fisher 18090010). Processed samples were then subjected to RT-qPCR analysis using primer pairs targeting *Ep400* (Table S1). To analyze protein expression, cells were harvested and subjected to western blot according to the conditions described above. To enable estimation of residual EP400 protein levels, a serial dilution of control (siNT-treated) sample was run alongside experimental samples.

#### BRM014 Treatment and ATAC Library Preparation

Fresh media containing 1 µM BRM014 was provided 72 h after the initial transfection, and cells were harvested after an additional 4 h of inhibitor treatment (for a total time 76 h between transfection and harvest). Cells were observed regularly to ensure that no large-scale defects in growth or viability occurred under these treatment conditions. After harvest, ATAC libraries were prepared according to the protocol detailed above. Libraries were sequenced at The Bauer Core Facility at Harvard University on an Illumina NovaSeq using an S4 flow cell and a paired-end 100-bp cycle run

### RNA-seq Library Preparation

H1299 and A549 cells were treated in triplicate with 5 µM BRM014 for 12 hours. Cells were harvested and resuspended in 500 µL TRIzol. To each sample an equal amount of the ERCC spike-in was added per cell to allow absolute quantification. RNA was extracted by chloroform precipitation and DNase (Invitrogen DNase I 18068015) treated. 500 ng of total RNA was used to make libraries with the TruSeq Stranded Total RNA Gold sequencing kit (Illumina 20020598). Two modifications to the manufacturer’s protocol were made. First, Superscript III was used rather than SuperScript II for the reverse transcription. Second, the A549 and H1299 samples were subject to 9 and 8 cycles of PCR amplification, respectively. Libraries were sequenced at The Bauer Core Facility at Harvard University on an Illumina NovaSeq using an SP flow cell and a paired-end 100-bp cycle run.

### Generation of *EP400*-KO A549 cells

Alt-R CRISPR-Cas9 crRNA targeting EP400 was ordered from IDT and annealed with ATTO550-labelled Alt-R tracrRNA (IDT 1072533), in an equimolar mixture at a final concentration of 100 µM. 1 µL of annealed RNAs was incubated with 1 µL of 10 mg/mL Cas9 protein (PNA Bio # CP01-200) at room temperature (22°C) for 25 minutes. The resultant riboprotein complex was introduced into cells by nucleofection (4D-Nucleofector X unit, Lonza bioscience), using the SF cell line kit and A549 cell program (CM 130). Two days after nucleofection, single cells positive for ATTO550 were isolated by fluorescence-activated cell sorting (FACS). After expanding single cell clones, homozygous disruption of *EP400* was confirmed by PCR of genomic DNA flanking the Cas9 cut site and Sanger sequencing. Sanger sequencing traces were compared using Inference of CRISPR Edits (ICE).^87^ Wildtype and clonal cell lines were then interrogated for *EP400* expression. To analyze mRNA expression, RNA was extracted using the Direct-zol RNA miniprep kit (VWR 76211-340). cDNA synthesis was performed using hexamer primers and SuperScript IV Reverse Transcriptase (Thermo Fisher 18090010). Processed samples were then subjected to RT-qPCR analysis using primer pairs targeting *EP400* and *ACTB* for normalization.

### Cell Proliferation Assay

Wildtype and *EP400-*KO A549 cells were plated at 300 cells per well in 100 µL media in 96 well plates. The following day, cells were treated with BRM014 or AU-15330 and then assayed after 8 days. Cell growth was determined using Cell Titer-Glo 2.0 Cell Viability Assay (Promega G9242). IC50 values were calculated in Prism using a four parameter non-linear fit inhibitor vs response model.

## QUANTIFICATION AND STATISTICAL ANALYSIS

### ATAC-seq data processing and mapping

All custom scripts described here are accessible at zenodo.^77^ Cutadapt 1.14^78^ was used to trim paired-end reads to 40 bp to remove adaptor sequences and low-quality reads. In order to identify spike-in reads, read pairs were next aligned to the *D. melanogaster* genome (dm6) using bowtie 1.2.2 (-k1 -v2 -X1000, —best -3 1 -p 5 —allow-contain —un).^72^ All reads that failed to align to the spike genome were subsequently aligned to the *M. musculus* genome (mm10) using bowtie 1.2.2 (-k1 -v2 -X1000 —best -3 1 -p 5 -S —allow-contain). The markdup tool (samtools 1.9)^80^ was used to flag duplicate reads, which were then discarded. Fragments were filtered to retain unique reads between 10 and 150 bp, representing regions of accessible chromatin, which were then converted to bedGraph format using the custom script extract_fragments.pl. As the replicate samples were highly correlated across ATAC-seq peaks and spike-in return rates were generally consistent across mESC samples, biological replicates were merged and depth-normalized using the custom scripts bedgraphs2stdBedGraph.pl and normalize_bedGraph.pl. Data was binned in 50 bp windows to generate bedGraph files for UCSC Genome Browser visualization and downstream analysis. Mapped reads and Spearman correlations between replicates are shown below:

| Sample | Total reads | Number of mapped reads | Replicate Spearman correlations (reads summed +/- 300 bp from peak centers) |
| --- | --- | --- | --- |
| mESC WT ATAC-seq rep1 | 62697825 | 40927157 | 0.94-0.96 |
| mESC WT ATAC-seq rep2 | 58980583 | 36080172 | 0.94-0.96 |
| mESC WT ATAC-seq rep3 | 168574624 | 100198015 | 0.94-0.96 |
| mESC WT ATAC-seq rep4 | 46502293 | 27779825 | 0.94-0.96 |
| mESC DMSO 2h ATAC-seq rep1 | 130212716 | 64415064 | 0.92-0.93 |
| mESC DMSO 2h ATAC-seq rep2 | 157163723 | 92364211 | 0.92-0.93 |
| mESC DMSO 2h ATAC-seq rep3 | 101477699 | 48635293 | 0.92-0.93 |
| mESC BRM014 2h ATAC-seq rep1 | 69377420 | 42103200 | 0.94-0.95 |
| mESC BRM014 2h ATAC-seq rep2 | 56773634 | 29270581 | 0.94-0.95 |
| mESC BRM014 2h ATAC-seq rep3 | 10070588 | 5522290 | 0.94-0.95 |
| mESC DMSO 4h ATAC-seq rep1 | 31363319 | 17925819 | 0.94 |
| mESC DMSO 4h ATAC-seq rep2 | 40843256 | 18687002 | 0.94 |
| mESC BRM014 4h ATAC-seq rep1 | 16319382 | 8573425 | 0.94 |
| mESC BRM014 4h ATAC-seq rep2 | 28491265 | 16413884 | 0.93 |
| mESC DMSO 8h ATAC-seq rep1 | 19153700 | 9695625 | 0.92-0.94 |
| mESC DMSO 8h ATAC-seq rep2 | 29033108 | 12731907 | 0.92-0.94 |
| mESC DMSO 8h ATAC-seq rep3 | 30886422 | 13344105 | 0.92-0.94 |
| mESC BRM014 8h ATAC-seq rep1 | 24320127 | 11377650 | 0.94-0.94 |
| mESC BRM014 8h ATAC-seq rep2 | 34579052 | 17553798 | 0.94-0.94 |
| mESC BRM014 8h ATAC-seq rep3 | 62829283 | 35207217 | 0.94-0.94 |
| mESC siNT DMSO 4h ATAC-seq rep1 | 32310317 | 24645541 | 0.95 |
| mESC siNT DMSO 4h ATAC-seq, rep2 | 29360108 | 22036024 | 0.95 |
| mESC siNT BRM014 4h ATAC-seq rep1 | 29177680 | 13353373 | 0.94 |
| mESC siNT BRM014 4h ATAC-seq rep2 | 43344493 | 32931732 | 0.94 |
| mESC siEP400 DMSO 4h ATAC-seq rep1 | 31235724 | 22889689 | 0.93-0.95 |
| mESC siEP400 DMSO 4h ATAC-seq rep2 | 27756232 | 20984256 | 0.93-0.95 |
| mESC siEP400 DMSO 4h ATAC-seq rep3 | 27405134 | 20467472 | 0.93-0.95 |
| mESC siEP400 BRM014 4h ATAC-seq rep1 | 25189856 | 18890986 | 0.92-0.95 |
| mESC siEP400 BRM014 4h ATAC-seq rep2 | 36974588 | 27560990 | 0.92-0.95 |
| mESC siEP400 BRM014 4h ATAC-seq rep3 | 26767737 | 18633718 | 0.92-0.95 |

### PRO-seq data processing and mapping

All custom scripts described here are accessible at zenodo.^77^ The custom script trim_and_filter_PE.pl was used to trim FASTQ files to 41 bp and remove read pairs with minimum average base quality scores below 20. Subsequent removal of adaptor sequences and low-quality reads was performed using cutadapt 1.14, and any reads shorter than 20 nt were discarded. The 3’-most nucleotide was removed from each trimmed read, after which bowtie (1.2.2) was used to map reads to the *Drosophila* dm6 genome (-k1 -v2 -best -X100 –un) and determine spike return across samples. Unaligned reads were mapped to the mm10 reference genome using the same parameters. Uniquely aligned read pairs were separated, and the custom script bowtie2stdBedGraph.pl was used to generate single-nucleotide resolution bedGraph files based on 3’ end mapping positions. Biological replicates were depth normalized using the custom script normalize_bedGraph.pl. As biological replicates were highly correlated (as indicated in the table below) replicates were merged using the custom script bedgraphs2stdbedGraph.pl, and data was binned in 50 bp windows.

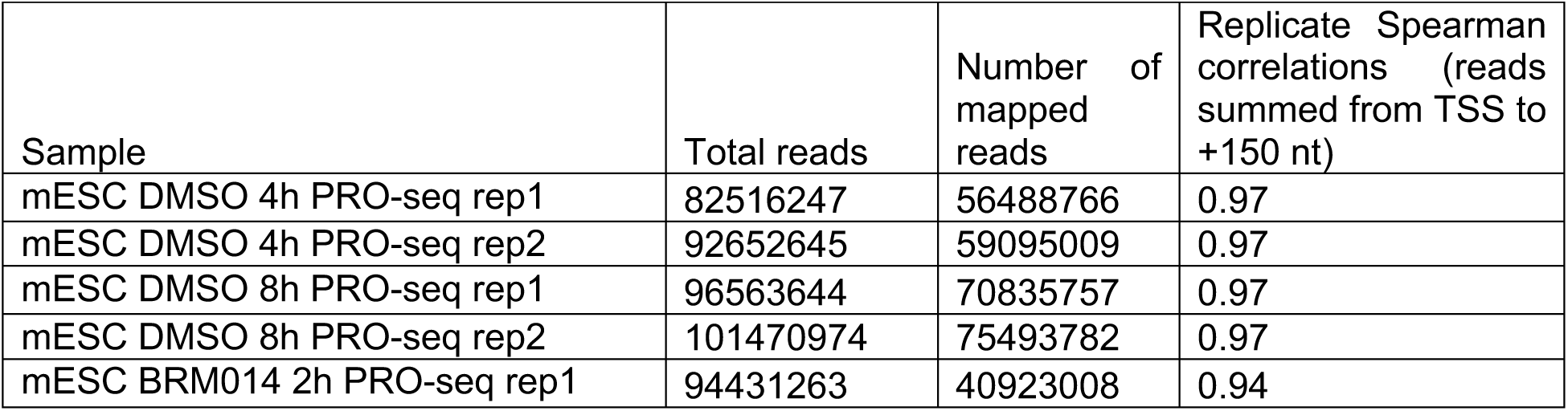
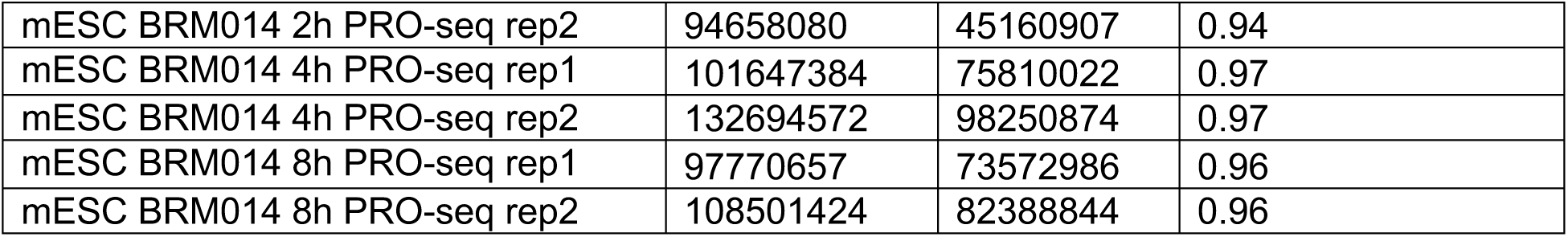

### Genome annotation

#### Transcription start sites

Genome-wide annotation of active transcripts and associated dominant transcription start site (TSS) and transcription end site (TES) locations was performed using the publicly available GetGeneAnnotations (GGA) pipeline.^76^ Briefly, GGA uses the 5’ end of PRO-seq reads to call TSSs and assign the dominant TSS for each gene. RNA-seq transcript isoform expression, quantified by kallisto (version 0.45.1),^81^ is then used to identify the most commonly used TES for each gene. GGA also enables comprehensive annotation of non-dominant TSSs and divergent obsTSSs (uaTSSs) associated with expressed genes. For this analysis, a total of 18,339 dominant and 1,671 non-dominant TSSs (and their associated TESs) were defined by GGA.

### RNA Biotype Analysis

Biotypes were derived from Ensembl annotations for mouse assembly GRCm38,p6 (v102).^88^ Promoters associated with biotypes in the “protein coding” category were designated as mRNA genes. Promoters associated with biotypes in the “long noncoding” and “short noncoding” categories were designated as ncRNA genes. Promoters associated with biotypes in the “pseudogene” category were designated as pseudogenes.

#### Candidate enhancer identification

To identify putative regulatory elements, peaks of bidirectional transcription were called from PRO-seq data using the dREG analysis tool^79^ under default parameters, generating a list of significant peaks (FDR < 0.05) with associated dREG scores, p-values, and peak center coordinates. Gene-distal elements (greater than 1.5 kb from a gene TSS) were retained as putative enhancers for downstream analysis. Peaks were subsequently filtered by read count, with peaks required to contain a minimum of 30 PRO-seq reads, giving rise to a final list of 71,330 dREG-identified candidate enhancer peaks.

### Peak calling and filtering

Final bed files from untreated mESC ATAC-seq libraries (N=4) were merged for peak-calling with HOMER (4.10.3) findPeaks using the “-style factor” argument.^82^ An initial list of 141,175 peaks was generated by this analysis, which was then filtered by peak score > 4.5 to generate a final list of 83,201 peaks. The HOMER annotatePeaks.pl command was used to associate each peak with nearby TSSs, as defined above. Peaks that were located within 1.5 kb of a dominant TSS (*n* = 17,160) were classified as proximal and shifted to center the associated TSS before subsequent analysis. After removal of duplicate TSSs, a final list of 13,536 sites was produced. For clarity, peaks that were located within 1.5 kb of a non-dominant TSS (*n* = 768) were excluded from further analysis. Remaining peaks were classified as distal (*n* = 65,273). Peaks that were located within 500 bp of a dREG-identified candidate enhancer were classified as enhancers (*n* = 32,149) and retained for analysis. To facilitate analysis of promoter-enhancer coordination, the HOMER annotatePeaks.pl command was also used to associate each promoter with its nearest enhancer and vice versa.

### ChIP-seq data processing and mapping

All custom scripts described here are accessible at zenodo.^77^ Adapter sequences were trimmed using cutadapt 1.14. Reads were first aligned to the *Drosophila* dm6 genome using bowtie 1.2.2, after which unaligned reads were mapped to the mm10 reference genome using analogous parameters. The custom script extract_fragments.pl was used to generate a final bedGraph file for each sample using uniquely mapped reads between 50 and 500 bp. BRM014- and DMSO-treated samples differed significantly in terms of spike-in read return. Therefore, the custom script normalize_bedGraph.pl was used to normalize individual libraries. As biological replicates were highly correlated (as indicated in the table below) replicates were merged using the custom script bedgraphs2stdbedGraph.pl, and data was binned in 50 bp windows.

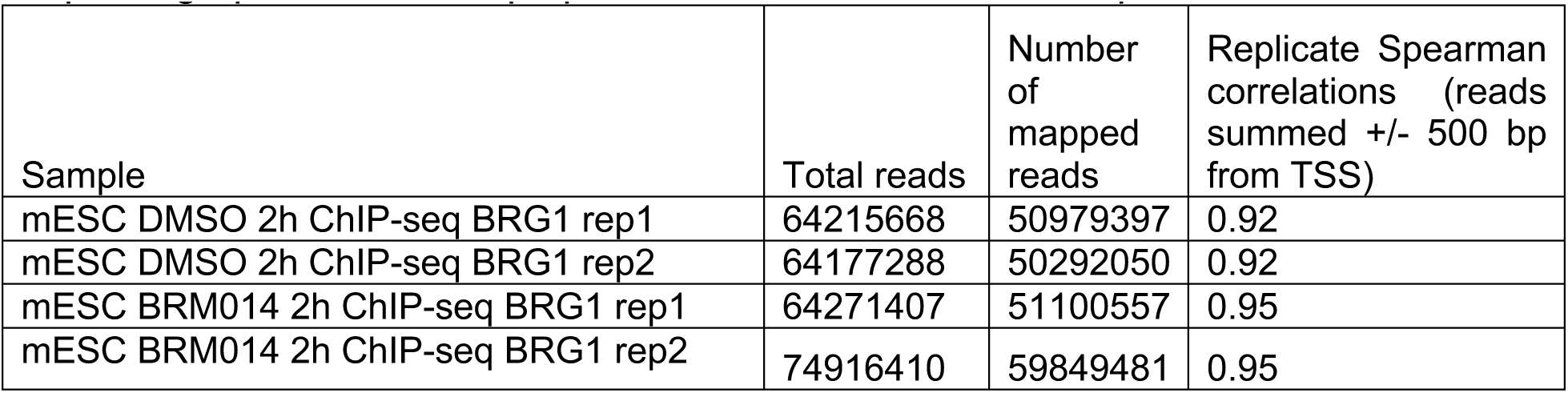

| Sample | Total reads | Number of mapped reads | Replicate Spearman correlations (reads summed +/- 500 bp from TSS) |
| --- | --- | --- | --- |
| mESC DMSO 2h ChIP-seq BRG1 rep1 | 64215668 | 50979397 | 0.92 |
| mESC DMSO 2h ChIP-seq BRG1 rep2 | 64177288 | 50292050 | 0.92 |
| mESC BRM014 2h ChIP-seq BRG1 rep1 | 64271407 | 51100557 | 0.95 |
| mESC BRM014 2h ChIP-seq BRG1 rep2 | 74916410 | 59849481 | 0.95 |

### Genome browser images

All genome browser images were generated from the UCSC Genome Browser (http://genome.ucsc.edu/)89 using genome build GRCm38/mm10.

### Metagenes and heatmaps

Composite metagene plots were generated by summing reads in 50 bp/nt bins at each indicated position relative to the TSS (promoters) or peak center (enhancers) using the custom script make_heatmap.pl,^77^ then dividing by the total number of sites. For PRO-seq data, 17 rRNA loci with aberrantly high signal were removed before final composite metagene plots were generated. Heatmaps were generated using Partek Genomics Suite (version 6.16.0812) from matrices summing reads in 50 bp/nt bins +/- 2 kb relative to the TSS (promoters) or peak center (enhancers). ΔATAC-seq and ΔPRO-seq heatmaps were generated by subtracting DMSO matrix values from the matrix values of the associated BRM014-treated sample, such that negative values correspond to regions of reduced signal following BRM014 treatment, and positive values correspond to regions of increased signal following BRM014 treatment. To order heatmaps, ATAC-seq signal from BRM014- and DMSO-treated samples was summed over a 600 bp window (-450 to +149 relative to TSS for promoters, -300 to +299 relative to peak center for enhancers). The raw difference in signal (# reads BRM014 - # reads DMSO) was calculated for each site at each time point, and sites were ranked in ascending order such that sites with the largest losses of signal are oriented at the top of the heatmap, and sites with the largest gains of signal are oriented at the bottom of the heatmap. For plot of relative BRG1 signal at promoters, BRG1 ChIP-seq reads for each promoter were summed from -750 to +149 relative to the TSS. For plot of relative BRG1 signal at enhancers, BRG1 ChIP-seq reads for each enhancer were summed from +/- 500 relative to the peak center. With sites ranked by difference in ATAC-seq signal after 2 h BRM014, pruning was performed in Prism 8 (8.4.3) to report average values over 10 rows. Data was smoothed across adjacent bins, and minimum and maximum values were used to normalize values across a range of 0 to 1.

### Clustering

Partek Genomics Suite (6.16.0812) was used to perform partitioning (k-means) clustering on all promoters (*n* = 13,536) based on relative ATAC-seq signal (normalized to DMSO control) across the BRM014 treatment time course (summed over a 600 bp window from -450 to +149 relative to the TSS). This analysis defined four promoter clusters (designated as Clusters 1-4) for downstream analysis. Heatmaps of relative ATAC-seq and PRO-seq signal by cluster were generated based on relative signal over the windows described above. Values were log2-transformed, and sites were ordered based on cluster assignment as indicated. Sites within each cluster were unranked.

### Relative Accessibility Analysis

Relative accessibility was calculated as the ratio of ATAC-seq signal in BRM014-treated samples compared to matched DMSO controls. For promoters, signal was summed from -450 to +149 bp relative to the TSS. For enhancers, signal was summed from -300 to +299 bp relative to the enhancer peak center. Values were log2-transformed before plotting.

### Relative PRO-seq Analysis

Relative promoter-proximal PRO-seq was calculated as the ratio of sense-strand PRO-seq signal from the TSS to +149 nt in BRM014-treated samples compared to matched DMSO controls. Relative gene-body PRO-seq signal was calculated as the ratio of sense-strand PRO-seq signal from +250 nt downstream of the TSS until one of the following conditions was met: (1) 500 bp upstream of the nearest enhancer; (2) the TES (as defined by GGA); or (3) a maximum of 5 kb. Relative enhancer PRO-seq signal was calculated as the ratio of PRO-seq signal on both strands in a window of -300 bp to +299 bp relative to the enhancer peak center. All values were log2-transformed before plotting.

### Differential Gene Expression Analysis

For each condition, PRO-seq 3’ read positions around each gene were counted from the dominant TSS+250 to the dominant TES. Sense PRO-seq reads were then counted for each gene and used as input for differential gene expression analysis, using DEseq2 to compare counts from each BRM014 treatment timepoint to those of matched DMSO controls. Up- and down-regulated genes were defined as genes exhibiting an increase or decrease of greater than 1.5-fold with BRM014 treatment, with an adjusted *P* value < 0.001.

### Gene Ontology Analysis

Gene ontology analysis was performed using the Database for Annotation, Visualization and Integrated Discovery (DAVID) v6.8 (https://david.ncifcrf.gov/home.jsp) under default parameters.^83, 84^ Cluster 2 was designated as background, and Cluster 1 was input as a gene list for analysis.

### Analysis of Publicly Available mESC Data

Previously published MNase-seq data^28^ were downloaded from the NCBI Gene Expression Omnibus as GSE85191 and aligned to mm10 according to the parameters described in the original work. MNase-seq data from control cells and cells treated with BRM014 for 24 h were downloaded as normalized wig files (GSE158345).^21^ Replicates were merged and converted to bedGraph format for metagene analysis. ATAC-seq data from control cells and BRG1-KO cells^8^ were downloaded (GSE87822) as FASTQ files and mapped according to the parameters described above. Promoter accessibility was calculated by summing signal from -450 to +149 bp relative to the TSS, and relative accessibility was calculated as the ratio of signal in BRG1 KO samples vs. control. Values were log2-transformed before plotting. TT-seq data^90^ were downloaded (GSE178230) and processed as described for PRO-seq data through mapping to the spike genome, after which STAR (v. 2.7.3a)^73^ was used to align data to the mm10 mouse genome. H3K4me3 ChIP-seq data were downloaded from the same source and processed as described above. Associated H3K27ac ChIP-seq data were retrieved through the 4DN Data Portal (https://data.4dnucleome.org/) at accession no. 4DNESQ33L4G7. Published H3K4me1 ChIP-seq data^61^ were downloaded from GSE56098. ChIP-seq data for CHD1, CHD2, CHD4, and EP400^3^ were downloaded from GSE64825. TIP60 ChIP-seq data^64^ were downloaded from GSE69671. SNF2H ChIP-seq data^63^ were downloaded from GSE123670. All samples were processed as described above. Processed data for CTCF ChIP-seq were downloaded from GSE137272. Processed data for OCT4, SOX2, and NANOG ChIP-seq were downloaded from GSE87822. CpG Island and GC Percent data tracks for the mm10 genome were downloaded from the UCSC Genome Browser Database as bedGraph files using the Table Browser tool.^62^ Previously published classifications were used to define bivalent genes,^91^ and the Ensembl BioMart^88^ was used to match Refseq and Ensembl gene IDs.

### Cancer Cell Line RNA-seq and ATAC-seq Analysis

#### RNA-seq analysis

RNA-seq FASTQ data files from DMSO and AU-15330-treated LNCaP and VCaP cells^25^ were downloaded from the sequence read archive (SRP313558). GetGeneAnnotations (GGA) scripts^76^ were used to annotate dominant active TSS and TES positions from LNCaP, VCaP, A549, and H1299 RNA-seq data. To quantify gene expression changes following BRM014 or AU-15330 treatment, RNA-seq samples were mapped to the hg38 genome using STAR version 2.7.3a.^73^ Gene counts were generated using featurecounts function of the Rsubread package version 2.0.1,^74^ and log2 fold change following BRM014 or AU-15330 treatment calculated with DESeq2 version 1.26.0.^75^ Protein-coding genes were filtered for a minimum of 0.3 FPKM counts in at least one condition and promoter ATAC-seq reads above the bottom 5^th^ percentile.

#### ATAC-seq Analysis

ATAC-seq FASTQ data files from control LNCaP (GSE171523^25^), VCaP (GSE171523^25^), A549 (GSE169955^69^), and H1299 (GSE141060^71^) cells were downloaded from the sequence read archive. ATAC-seq data were mapped to hg38 using the same parameters described above for ATAC-seq mapping in mESCs. ATAC-seq counts for each gene were summed in a window of -500 bp to +499 bp around the dominant TSS.

#### H3K4me1 ChIP-seq Analysis

H3K4me1 ChIP-seq FASTQ data files from control LNCaP (GSE122922^68^), VCaP (GSE148400^67^), A549 (GSE169955^69^), and H1299 (DRR016953^70^) cells were downloaded from the sequence read archive (see key resource table). ATAC-seq data were mapped to hg38 using the same parameters described above for mapping in mESCs. ATAC-seq counts for each gene were summed in a window of -500 bp to +499 bp around the dominant TSS.

#### Predicting genes sensitive or resistant to SWI/SNF inhibition

H3K4me1 and ATAC-seq were used to predict genes sensitive or resistant to SWI/SNF perturbation by BRM014 or AU-15330. Genes with promoter H3K4me1 in the top 15% and ATAC-seq in the bottom 15% were predicted to be sensitive. Genes with promoter H3K4me1 in the bottom 15% and ATAC-seq signal in the top 15% were predicted to be resistant. For ATAC-seq only predictions, genes in the bottom and top 5% ATAC-seq signal were predicted to be sensitive and resistant respectively. The number of genes predicted to be sensitive or resistant in each cell line is shown below:

| Cell line | Prediction | Number of genes |
| --- | --- | --- |
| A549 | Sensitive – H3K4me1 + ATAC-seq | 634 |
| H1299 | Sensitive – H3K4me1 + ATAC-seq | 204 |
| LNCaP | Sensitive – H3K4me1 + ATAC-seq | 364 |
| VCaP | Sensitive – H3K4me1 + ATAC-seq | 567 |
| A549 | Resistant – H3K4me1 + ATAC-seq | 370 |
| H1299 | Resistant – H3K4me1 + ATAC-seq | 327 |
| LNCaP | Resistant – H3K4me1 + ATAC-seq | 252 |
| VCaP | Resistant – H3K4me1 + ATAC-seq | 444 |
| A549 | Sensitive – ATAC-seq | 604 |
| H1299 | Sensitive – ATAC-seq | 593 |
| LNCaP | Sensitive – ATAC-seq | 677 |
| VCaP | Sensitive – ATAC-seq | 704 |
| A549 | Resistant – ATAC-seq | 598 |
| H1299 | Resistant – ATAC-seq | 593 |
| LNCaP | Resistant – ATAC-seq | 580 |
| VCaP | Resistant – ATAC-seq | 668 |

### Box plots and statistical analysis

Box plots were plotted in GraphPad Prism 9.0 (GraphPad Software, San Diego, CA, USA) and have a line at the median, and whiskers show the 10-90th percentiles. P-values were calculated in Prism, using the indicated statistical test, except for the overlap in Venn diagrams which were calculated using the phyper function in R (3.6.1).

## REFERENCES

1. Clapier, C.R., Iwasa, J., Cairns, B.R., and Peterson, C.L. (2017). Mechanisms of action and regulation of ATP-dependent chromatin-remodelling complexes. Nat Rev Mol Cell Bio 18, 407–422. 10.1038/nrm.2017.26.

2. Narlikar, G.J., Sundaramoorthy, R., and Owen-Hughes, T. (2013). Mechanisms and Functions of ATP-Dependent Chromatin-Remodeling Enzymes. Cell 154, 490–503. 10.1016/j.cell.2013.07.011.

3. Dieuleveult, M. de, Yen, K., Hmitou, I., Depaux, A., Boussouar, F., Dargham, D.B., Jounier, S., Humbertclaude, H., Ribierre, F., Baulard, C., et al. (2016). Genome-wide nucleosome specificity and function of chromatin remodellers in ES cells. Nature 530, 113–116. 10.1038/nature16505.

4. Barisic, D., Stadler, M.B., Iurlaro, M., and Schübeler, D. (2019). Mammalian ISWI and SWI/SNF selectively mediate binding of distinct transcription factors. Nature 569, 136–140. 10.1038/s41586-019-1115-5.

5. Brown, S.A., Imbalzano, A.N., and Kingston, R.E. (1996). Activator-dependent regulation of transcriptional pausing on nucleosomal templates. Gene Dev 10, 1479–1490. 10.1101/gad.10.12.1479.

6. Nakayama, R.T., Pulice, J.L., Valencia, A.M., McBride, M.J., McKenzie, Z.M., Gillespie, M.A., Ku, W.L., Teng, M., Cui, K., Williams, R.T., et al. (2017). SMARCB1 is required for widespread BAF complex–mediated activation of enhancers and bivalent promoters. Nat Genet 49, 1613–1623. 10.1038/ng.3958.

7. Alver, B.H., Kim, K.H., Lu, P., Wang, X., Manchester, H.E., Wang, W., Haswell, J.R., Park, P.J., and Roberts, C.W.M. (2017). The SWI/SNF chromatin remodelling complex is required for maintenance of lineage specific enhancers. Nat Commun 8, 14648. 10.1038/ncomms14648.

8. King, H.W., and Klose, R.J. (2017). The pioneer factor OCT4 requires the chromatin remodeller BRG1 to support gene regulatory element function in mouse embryonic stem cells. Elife 6, e22631. 10.7554/elife.22631.

9. Kadoch, C., Hargreaves, D.C., Hodges, C., Elias, L., Ho, L., Ranish, J., and Crabtree, G.R. (2013). Proteomic and bioinformatic analysis of mammalian SWI/SNF complexes identifies extensive roles in human malignancy. Nat Genet 45, 592–601. 10.1038/ng.2628.

10. Centore, R.C., Sandoval, G.J., Soares, L.M.M., Kadoch, C., and Chan, H.M. (2020). Mammalian SWI/SNF Chromatin Remodeling Complexes: Emerging Mechanisms and Therapeutic Strategies. Trends Genet 36, 936–950. 10.1016/j.tig.2020.07.011.

11. Hainer, S.J., Gu, W., Carone, B.R., Landry, B.D., Rando, O.J., Mello, C.C., and Fazzio, T.G. (2015). Suppression of pervasive noncoding transcription in embryonic stem cells by esBAF. Gene Dev 29, 362–378. 10.1101/gad.253534.114.

12. Tolstorukov, M.Y., Sansam, C.G., Lu, P., Koellhoffer, E.C., Helming, K.C., Alver, B.H., Tillman, E.J., Evans, J.A., Wilson, B.G., Park, P.J., et al. (2013). Swi/Snf chromatin remodeling/tumor suppressor complex establishes nucleosome occupancy at target promoters. Proc National Acad Sci 110, 10165–10170. 10.1073/pnas.1302209110.

13. Hodges, H.C., Stanton, B.Z., Cermakova, K., Chang, C.-Y., Miller, E.L., Kirkland, J.G., Ku, W.L., Veverka, V., Zhao, K., and Crabtree, G.R. (2018). Dominant-negative SMARCA4 mutants alter the accessibility landscape of tissue-unrestricted enhancers. Nat Struct Mol Biol 25, 61–72. 10.1038/s41594-017-0007-3.

14. Flynn, R.A., Do, B.T., Rubin, A.J., Calo, E., Lee, B., Kuchelmeister, H., Rale, M., Chu, C., Kool, E.T., Wysocka, J., et al. (2016). 7SK-BAF axis controls pervasive transcription at enhancers. Nat Struct Mol Biol 23, 231–238. 10.1038/nsmb.3176.

15. Weber, C.M., Hafner, A., Kirkland, J.G., Braun, S.M.G., Stanton, B.Z., Boettiger, A.N., and Crabtree, G.R. (2021). mSWI/SNF promotes Polycomb repression both directly and through genome-wide redistribution. Nat Struct Mol Biol 28, 501–511. 10.1038/s41594-021-00604-7.

16. Zhang, X., Li, B., Li, W., Ma, L., Zheng, D., Li, L., Yang, W., Chu, M., Chen, W., Mailman, R.B., et al. (2014). Transcriptional Repression by the BRG1-SWI/SNF Complex Affects the Pluripotency of Human Embryonic Stem Cells. Stem Cell Rep 3, 460–474. 10.1016/j.stemcr.2014.07.004.

17. Bossen, C., Murre, C.S., Chang, A.N., Mansson, R., Rodewald, H.-R., and Murre, C. (2015). The chromatin remodeler Brg1 activates enhancer repertoires to establish B cell identity and modulate cell growth. Nat Immunol 16, 775–784. 10.1038/ni.3170.

18. Park, Y.-K., Lee, J.-E., Yan, Z., McKernan, K., O’Haren, T., Wang, W., Peng, W., and Ge, K. (2021). Interplay of BAF and MLL4 promotes cell type-specific enhancer activation. Nat Commun 12, 1630. 10.1038/s41467-021-21893-y.

19. Papillon, J.P.N., Nakajima, K., Adair, C.D., Hempel, J., Jouk, A.O., Karki, R.G., Mathieu, S., Möbitz, H., Ntaganda, R., Smith, T., et al. (2018). Discovery of Orally Active Inhibitors of Brahma Homolog (BRM)/SMARCA2 ATPase Activity for the Treatment of Brahma Related Gene 1 (BRG1)/SMARCA4-Mutant Cancers. J Med Chem 61, 10155–10172. 10.1021/acs.jmedchem.8b01318.

20. Schick, S., Grosche, S., Kohl, K.E., Drpic, D., Jaeger, M.G., Marella, N.C., Imrichova, H., Lin, J.-M.G., Hofstätter, G., Schuster, M., et al. (2021). Acute BAF perturbation causes immediate changes in chromatin accessibility. Nat Genet, 1–10. 10.1038/s41588-021-00777-3.

21. Iurlaro, M., Stadler, M.B., Masoni, F., Jagani, Z., Galli, G.G., and Schübeler, D. (2021). Mammalian SWI/SNF continuously restores local accessibility to chromatin. Nat Genet, 1–9. 10.1038/s41588-020-00768-w.

22. Biggar, S.R., and Crabtree, G.R. (1999). Continuous and widespread roles for the Swi–Snf complex in transcription. Embo J 18, 2254–2264. 10.1093/emboj/18.8.2254.

23. Schwabish, M.A., and Struhl, K. (2007). The Swi/Snf Complex Is Important for Histone Eviction during Transcriptional Activation and RNA Polymerase II Elongation In Vivo. Mol Cell Biol 27, 6987–6995. 10.1128/mcb.00717-07.

24. Sudarsanam, P., Cao, Y., Wu, L., Laurent, B.C., and Winston, F. (1999). The nucleosome remodeling complex, Snf/Swi, is required for the maintenance of transcription in vivo and is partially redundant with the histone acetyltransferase, Gcn5. Embo J 18, 3101–3106. 10.1093/emboj/18.11.3101.

25. Xiao, L., Parolia, A., Qiao, Y., Bawa, P., Eyunni, S., Mannan, R., Carson, S.E., Chang, Y., Wang, X., Zhang, Y., et al. (2022). Targeting SWI/SNF ATPases in enhancer-addicted prostate cancer. Nature 601, 434–439. 10.1038/s41586-021-04246-z.

26. Jagani, Z., Chenail, G., Xiang, K., Bushold, G., Bhang, H.-E.C., Li, A., Elliott, G., Zhu, J., Vattay, A., Gilbert, T., et al. (2019). In-Depth Characterization and Validation in BRG1-Mutant Lung Cancers Define Novel Catalytic Inhibitors of SWI/SNF Chromatin Remodeling. Biorxiv, 812628. 10.1101/812628.

27. Mahat, D.B., Kwak, H., Booth, G.T., Jonkers, I.H., Danko, C.G., Patel, R.K., Waters, C.T., Munson, K., Core, L.J., and Lis, J.T. (2016). Base-pair-resolution genome-wide mapping of active RNA polymerases using precision nuclear run-on (PRO-seq). Nat Protoc 11, 1455– 1476. 10.1038/nprot.2016.086.

28. Henriques, T., Scruggs, B.S., Inouye, M.O., Muse, G.W., Williams, L.H., Burkholder, A.B., Lavender, C.A., Fargo, D.C., and Adelman, K. (2018). Widespread transcriptional pausing and elongation control at enhancers. Gene Dev 32, 26–41. 10.1101/gad.309351.117.

29. Tippens, N.D., Liang, J., Leung, A.K.-Y., Wierbowski, S.D., Ozer, A., Booth, J.G., Lis, J.T., and Yu, H. (2020). Transcription imparts architecture, function, and logic to enhancer units. Nat Genet 52, 1067–1075. 10.1038/s41588-020-0686-2.

30. Wang, Z., Chivu, A.G., Choate, L.A., Rice, E.J., Miller, D.C., Chu, T., Chou, S.-P., Kingsley, N.B., Petersen, J.L., Finno, C.J., et al. (2022). Prediction of histone post-translational modification patterns based on nascent transcription data. Nat Genet 54, 295–305. 10.1038/s41588-022-01026-x.

31. Danko, C.G., Hyland, S.L., Core, L.J., Martins, A.L., Waters, C.T., Lee, H.W., Cheung, V.G., Kraus, W.L., Lis, J.T., and Siepel, A. (2015). Identification of active transcriptional regulatory elements with GRO-seq. Nat Methods 12, 433–438. 10.1038/nmeth.3329.

32. Ho, L., Ronan, J.L., Wu, J., Staahl, B.T., Chen, L., Kuo, A., Lessard, J., Nesvizhskii, A.I., Ranish, J., and Crabtree, G.R. (2009). An embryonic stem cell chromatin remodeling complex, esBAF, is essential for embryonic stem cell self-renewal and pluripotency. Proc National Acad Sci 106, 5181–5186. 10.1073/pnas.0812889106.

33. Bartholomew, B. (2014). Regulating the Chromatin Landscape: Structural and Mechanistic Perspectives. Biochemistry-us 83, 671–696. 10.1146/annurev-biochem-051810-093157.

34. Tilly, B.C., Chalkley, G.E., Knaap, J.A. van der, Moshkin, Y.M., Kan, T.W., Dekkers, D.H., Demmers, J.A., and Verrijzer, C.P. (2021). In vivo analysis reveals that ATP-hydrolysis couples remodeling to SWI/SNF release from chromatin. Elife 10, e69424. 10.7554/elife.69424.

35. Wiechens, N., Singh, V., Gkikopoulos, T., Schofield, P., Rocha, S., and Owen-Hughes, T. (2016). The Chromatin Remodelling Enzymes SNF2H and SNF2L Position Nucleosomes adjacent to CTCF and Other Transcription Factors. Plos Genet 12, e1005940. 10.1371/journal.pgen.1005940.

36. Bomber, M.L., Wang, J., Liu, Q., Barnett, K.R., Layden, H.M., Hodges, E., Stengel, K.R., and Hiebert, S.W. (2023). Human SMARCA5 is continuously required to maintain nucleosome spacing. Mol Cell. 10.1016/j.molcel.2022.12.018.

37. Engreitz, J.M., Haines, J.E., Perez, E.M., Munson, G., Chen, J., Kane, M., McDonel, P.E., Guttman, M., and Lander, E.S. (2016). Local regulation of gene expression by lncRNA promoters, transcription and splicing. Nature 539, 452–455. 10.1038/nature20149.

38. Hota, S.K., Johnson, J.R., Verschueren, E., Thomas, R., Blotnick, A.M., Zhu, Y., Sun, X., Pennacchio, L.A., Krogan, N.J., and Bruneau, B.G. (2019). Dynamic BAF chromatin remodeling complex subunit inclusion promotes temporally distinct gene expression programs in cardiogenesis. Development 146, dev174086. 10.1242/dev.174086.

39. Matsumoto, S., Banine, F., Struve, J., Xing, R., Adams, C., Liu, Y., Metzger, D., Chambon, P., Rao, M.S., and Sherman, L.S. (2006). Brg1 is required for murine neural stem cell maintenance and gliogenesis. Dev Biol 289, 372–383. 10.1016/j.ydbio.2005.10.044.

40. Lessard, J., Wu, J.I., Ranish, J.A., Wan, M., Winslow, M.M., Staahl, B.T., Wu, H., Aebersold, R., Graef, I.A., and Crabtree, G.R. (2007). An Essential Switch in Subunit Composition of a Chromatin Remodeling Complex during Neural Development. Neuron 55, 201–215. 10.1016/j.neuron.2007.06.019.

41. Ramirez-Carrozzi, V.R., Braas, D., Bhatt, D.M., Cheng, C.S., Hong, C., Doty, K.R., Black, J.C., Hoffmann, A., Carey, M., and Smale, S.T. (2009). A Unifying Model for the Selective Regulation of Inducible Transcription by CpG Islands and Nucleosome Remodeling. Cell 138, 114–128. 10.1016/j.cell.2009.04.020.

42. Fenouil, R., Cauchy, P., Koch, F., Descostes, N., Cabeza, J.Z., Innocenti, C., Ferrier, P., Spicuglia, S., Gut, M., Gut, I., et al. (2012). CpG islands and GC content dictate nucleosome depletion in a transcription-independent manner at mammalian promoters. Genome Res 22, 2399–2408. 10.1101/gr.138776.112.

43. Core, L.J., Martins, A.L., Danko, C.G., Waters, C.T., Siepel, A., and Lis, J.T. (2014). Analysis of nascent RNA identifies a unified architecture of initiation regions at mammalian promoters and enhancers. Nat Genet 46, 1311–1320. 10.1038/ng.3142.

44. Oike, T., Ogiwara, H., Tominaga, Y., Ito, K., Ando, O., Tsuta, K., Mizukami, T., Shimada, Y., Isomura, H., Komachi, M., et al. (2013). A Synthetic Lethality–Based Strategy to Treat Cancers Harboring a Genetic Deficiency in the Chromatin Remodeling Factor BRG1. Cancer Res 73, 5508–5518. 10.1158/0008-5472.can-12-4593.

45. Vangamudi, B., Paul, T.A., Shah, P.K., Kost-Alimova, M., Nottebaum, L., Shi, X., Zhan, Y., Leo, E., Mahadeshwar, H.S., Protopopov, A., et al. (2015). The SMARCA2/4 ATPase Domain Surpasses the Bromodomain as a Drug Target in SWI/SNF-Mutant Cancers: Insights from cDNA Rescue and PFI-3 Inhibitor Studies. Cancer Res 75, 3865–3878. 10.1158/0008-5472.can-14-3798.

46. Doyon, Y., Selleck, W., Lane, W.S., Tan, S., and Côté, J. (2004). Structural and Functional Conservation of the NuA4 Histone Acetyltransferase Complex from Yeast to Humans. Mol Cell Biol 24, 1884–1896. 10.1128/mcb.24.5.1884-1896.2004.

47. Wichmann, J., Pitt, C., Eccles, S., Garnham, A.L., Li-Wai-Suen, C.S.N., May, R., Allan, E., Wilcox, S., Herold, M.J., Smyth, G.K., et al. (2022). Loss of TIP60 (KAT5) abolishes H2AZ lysine 7 acetylation and causes p53, INK4A, and ARF-independent cell cycle arrest. Cell Death Dis 13, 627. 10.1038/s41419-022-05055-6.

48. Kim, S., Natesan, S., Cornilescu, G., Carlson, S., Tonelli, M., McClurg, U.L., Binda, O., Robson, C.N., Markley, J.L., Balaz, S., et al. (2016). Mechanism of Histone H3K4me3 Recognition by the Plant Homeodomain of Inhibitor of Growth 3*. J Biol Chem 291, 18326– 18341. 10.1074/jbc.m115.690651.

49. Shi, X., Hong, T., Walter, K.L., Ewalt, M., Michishita, E., Hung, T., Carney, D., Peña, P., Lan, F., Kaadige, M.R., et al. (2006). ING2 PHD domain links histone H3 lysine 4 methylation to active gene repression. Nature 442, 96–99. 10.1038/nature04835.

50. Peña, P.V., Davrazou, F., Shi, X., Walter, K.L., Verkhusha, V.V., Gozani, O., Zhao, R., and Kutateladze, T.G. (2006). Molecular mechanism of histone H3K4me3 recognition by plant homeodomain of ING2. Nature 442, 100–103. 10.1038/nature04814.

51. Halaburkova, A., Cahais, V., Novoloaca, A., Araujo, M.G. da S., Khoueiry, R., Ghantous, A., and Herceg, Z. (2020). Pan-cancer multi-omics analysis and orthogonal experimental assessment of epigenetic driver genes. Genome Res 30, 1517–1532. 10.1101/gr.268292.120.

52. Fazzio, T.G., Huff, J.T., and Panning, B. (2008). An RNAi Screen of Chromatin Proteins Identifies Tip60-p400 as a Regulator of Embryonic Stem Cell Identity. Cell 134, 162–174. 10.1016/j.cell.2008.05.031.

53. Fazzio, T.G., Huff, J.T., and Panning, B. (2008). Chromatin regulation Tip(60)s the balance in embryonic stem cell self-renewal. Cell Cycle 7, 3302–3306. 10.4161/cc.7.21.6928.

54. Jin, C., Zang, C., Wei, G., Cui, K., Peng, W., Zhao, K., and Felsenfeld, G. (2009). H3.3/H2A.Z double variant–containing nucleosomes mark “nucleosome-free regions” of active promoters and other regulatory regions. Nat Genet 41, 941–945. 10.1038/ng.409.

55. Jin, C., and Felsenfeld, G. (2007). Nucleosome stability mediated by histone variants H3.3 and H2A.Z. Gene Dev 21, 1519–1529. 10.1101/gad.1547707.

56. Colino-Sanguino, Y., Clark, S.J., and Valdes-Mora, F. (2016). H2A.Z acetylation and transcription: ready, steady, go. Epigenomics-uk 8, 583–586. 10.2217/epi-2016-0016.

57. Halley, J.E., Kaplan, T., Wang, A.Y., Kobor, M.S., and Rine, J. (2010). Roles for H2A.Z and Its Acetylation in GAL1 Transcription and Gene Induction, but Not GAL1-Transcriptional Memory. Plos Biol 8, e1000401. 10.1371/journal.pbio.1000401.

58. Law, C., and Cheung, P. (2015). Expression of Non-acetylatable H2A.Z in Myoblast Cells Blocks Myoblast Differentiation through Disruption of MyoD Expression*. J Biol Chem 290, 13234–13249. 10.1074/jbc.m114.595462.

59. Janas, J.A., Zhang, L., Luu, J.H., Demeter, J., Meng, L., Marro, S.G., Mall, M., Mooney, N.A., Schaukowitch, K., Ng, Y.H., et al. (2022). Tip60-mediated H2A.Z acetylation promotes neuronal fate specification and bivalent gene activation. Mol Cell 82, 4627–4646.e14. 10.1016/j.molcel.2022.11.002.

60. Vlaming, H., Mimoso, C.A., Field, A.R., Martin, B.J.E., and Adelman, K. (2022). Screening thousands of transcribed coding and non-coding regions reveals sequence determinants of RNA polymerase II elongation potential. Nat Struct Mol Biol 29, 613–620. 10.1038/s41594-022-00785-9.

61. Buecker, C., Srinivasan, R., Wu, Z., Calo, E., Acampora, D., Faial, T., Simeone, A., Tan, M., Swigut, T., and Wysocka, J. (2014). Reorganization of Enhancer Patterns in Transition from Naive to Primed Pluripotency. Cell Stem Cell 14, 838–853. 10.1016/j.stem.2014.04.003.

62. Karolchik, D., Hinrichs, A.S., Furey, T.S., Roskin, K.M., Sugnet, C.W., Haussler, D., and Kent, W.J. (2004). The UCSC Table Browser data retrieval tool. Nucleic Acids Res 32, D493– D496. 10.1093/nar/gkh103.

63. Song, Y., Liang, Z., Zhang, J., Hu, G., Wang, J., Li, Y., Guo, R., Dong, X., Babarinde, I.A., Ping, W., et al. (2022). CTCF functions as an insulator for somatic genes and a chromatin remodeler for pluripotency genes during reprogramming. Cell Reports 39, 110626. 10.1016/j.celrep.2022.110626.

64. Ravens, S., Yu, C., Ye, T., Stierle, M., and Tora, L. (2015). Tip60 complex binds to active Pol II promoters and a subset of enhancers and co-regulates the c-Myc network in mouse embryonic stem cells. Epigenet Chromatin 8, 45. 10.1186/s13072-015-0039-z.

65. Hu, G., Cui, K., Northrup, D., Liu, C., Wang, C., Tang, Q., Ge, K., Levens, D., Crane-Robinson, C., and Zhao, K. (2013). H2A.Z Facilitates Access of Active and Repressive Complexes to Chromatin in Embryonic Stem Cell Self-Renewal and Differentiation. Cell Stem Cell 12, 180–192. 10.1016/j.stem.2012.11.003.

66. Justice, M., Carico, Z.M., Stefan, H.C., and Dowen, J.M. (2020). A WIZ/Cohesin/CTCF Complex Anchors DNA Loops to Define Gene Expression and Cell Identity. Cell Reports 31, 107503. 10.1016/j.celrep.2020.03.067.

67. Baumgart, S.J., Nevedomskaya, E., Lesche, R., Newman, R., Mumberg, D., and Haendler, B. (2020). Darolutamide antagonizes androgen signaling by blocking enhancer and super-enhancer activation. Mol Oncol 14, 2022–2039. 10.1002/1878-0261.12693.

68. Sugiura, M., Sato, H., Okabe, A., Fukuyo, M., Mano, Y., Shinohara, K., Rahmutulla, B., Higuchi, K., Maimaiti, M., Kanesaka, M., et al. (2020). Identification of AR-V7 downstream genes commonly targeted by AR/AR-V7 and specifically targeted by AR-V7 in castration resistant prostate cancer. Transl Oncol 14, 100915. 10.1016/j.tranon.2020.100915.

69. Dunham, I., Kundaje, A., Aldred, S.F., Collins, P.J., Davis, C.A., Doyle, F., Epstein, C.B., Frietze, S., Harrow, J., Kaul, R., et al. (2012). An integrated encyclopedia of DNA elements in the human genome. Nature 489, 57–74. 10.1038/nature11247.

70. Suzuki, A., Makinoshima, H., Wakaguri, H., Esumi, H., Sugano, S., Kohno, T., Tsuchihara, K., and Suzuki, Y. (2014). Aberrant transcriptional regulations in cancers: genome, transcriptome and epigenome analysis of lung adenocarcinoma cell lines. Nucleic Acids Res 42, 13557–13572. 10.1093/nar/gku885.

71. Kim, D., Kim, Y., Lee, B.B., Cho, E.Y., Han, J., Shim, Y.M., and Kim, D.-H. (2021). Metformin Reduces Histone H3K4me3 at the Promoter Regions of Positive Cell Cycle Regulatory Genes in Lung Cancer Cells. Cancers 13, 739. 10.3390/cancers13040739.

72. Langmead, B., Trapnell, C., Pop, M., and Salzberg, S.L. (2009). Ultrafast and memory-efficient alignment of short DNA sequences to the human genome. Genome Biol 10, R25. 10.1186/gb-2009-10-3-r25.

73. Dobin, A., Davis, C.A., Schlesinger, F., Drenkow, J., Zaleski, C., Jha, S., Batut, P., Chaisson, M., and Gingeras, T.R. (2013). STAR: ultrafast universal RNA-seq aligner. Bioinformatics 29, 15–21. 10.1093/bioinformatics/bts635.

74. Liao, Y., Smyth, G.K., and Shi, W. (2019). The R package Rsubread is easier, faster, cheaper and better for alignment and quantification of RNA sequencing reads. Nucleic Acids Res 47, gkz114-. 10.1093/nar/gkz114.

75. Love, M.I., Huber, W., and Anders, S. (2014). Moderated estimation of fold change and dispersion for RNA-seq data with DESeq2. Genome Biol 15, 550. 10.1186/s13059-014-0550-8.

76. Martin, B.J., Mimoso, C.A., and Adelman, K. (2021). AdelmanLab/GetGeneAnnotation_GGA: AdelmanLab/GetGeneAnnotation_GGA/v1. Zenodo. https://doi.org/10.5281/zenodo.5519928.

77. Martin, B.J., Adelman, K., and Nelson, G. (2021). AdelmanLab/NIH_scripts: AdelmanLab/NIH_scripts/v1. Zenodo. https://doi.org/10.5281/zenodo.5519915.

78. Martin, M. (2011). Cutadapt removes adapter sequence from high-throughput sequencing reads. EMBnet.journal 17, 10–12. https://doi.org/10.14806/ej.17.1.200.

79. Wang, Z., Chu, T., Choate, L.A., and Danko, C.G. (2019). Identification of regulatory elements from nascent transcription using dREG. Genome Res 29, 293–303. 10.1101/gr.238279.118.

80. Li, H., Handsaker, B., Wysoker, A., Fennell, T., Ruan, J., Homer, N., Marth, G., Abecasis, G., Durbin, R., and Subgroup, 1000 Genome Project Data Processing (2009). The Sequence Alignment/Map format and SAMtools. Bioinformatics 25, 2078–2079. 10.1093/bioinformatics/btp352.

81. Bray, N.L., Pimentel, H., Melsted, P., and Pachter, L. (2016). Near-optimal probabilistic RNA-seq quantification. Nat Biotechnol 34, 525–527. 10.1038/nbt.3519.

82. Heinz, S., Benner, C., Spann, N., Bertolino, E., Lin, Y.C., Laslo, P., Cheng, J.X., Murre, C., Singh, H., and Glass, C.K. (2010). Simple Combinations of Lineage-Determining Transcription Factors Prime cis-Regulatory Elements Required for Macrophage and B Cell Identities. Mol Cell 38, 576–589. 10.1016/j.molcel.2010.05.004.

83. Huang, D.W., Sherman, B.T., and Lempicki, R.A. (2009). Systematic and integrative analysis of large gene lists using DAVID bioinformatics resources. Nat Protoc 4, 44–57. 10.1038/nprot.2008.211.

84. Huang, D.W., Sherman, B.T., and Lempicki, R.A. (2009). Bioinformatics enrichment tools: paths toward the comprehensive functional analysis of large gene lists. Nucleic Acids Res 37, 1–13. 10.1093/nar/gkn923.

85. Buenrostro, J.D., Giresi, P.G., Zaba, L.C., Chang, H.Y., and Greenleaf, W.J. (2013). Transposition of native chromatin for fast and sensitive epigenomic profiling of open chromatin, DNA-binding proteins and nucleosome position. Nat Methods 10, 1213–1218. 10.1038/nmeth.2688.

86. Tian, B., Yang, J., and Brasier, A.R. (2011). Two-step cross-linking for analysis of protein-chromatin interactions. Methods Mol Biology 809, 105–120. 10.1007/978-1-61779-376-9_7.

87. Conant, D., Hsiau, T., Rossi, N., Oki, J., Maures, T., Waite, K., Yang, J., Joshi, S., Kelso, R., Holden, K., et al. (2022). Inference of CRISPR Edits from Sanger Trace Data. Crispr J 5, 123–130. 10.1089/crispr.2021.0113.

88. Yates, A.D., Achuthan, P., Akanni, W., Allen, J., Allen, J., Alvarez-Jarreta, J., Amode, M.R., Armean, I.M., Azov, A.G., Bennett, R., et al. (2020). Ensembl 2020. Nucleic Acids Res 48, D682–D688. 10.1093/nar/gkz966.

89. Kent, W.J., Sugnet, C.W., Furey, T.S., Roskin, K.M., Pringle, T.H., Zahler, A.M., and Haussler, and D. (2002). The Human Genome Browser at UCSC. Genome Res 12, 996–1006. 10.1101/gr.229102.

90. Vlaming, H., Mimoso, C.A., Martin, B.J., Field, A.R., and Adelman, K. (2021). Screening thousands of transcribed coding and non-coding regions reveals sequence determinants of RNA polymerase II elongation potential. Biorxiv, 2021.06.01.446655. 10.1101/2021.06.01.446655.

91. Ku, M., Koche, R.P., Rheinbay, E., Mendenhall, E.M., Endoh, M., Mikkelsen, T.S., Presser, A., Nusbaum, C., Xie, X., Chi, A.S., et al. (2008). Genomewide Analysis of PRC1 and PRC2 Occupancy Identifies Two Classes of Bivalent Domains. Plos Genet 4, e1000242. 10.1371/journal.pgen.1000242.

