## Supplemental material for "Global identification of direct SWI/SNF targets reveals compensation by EP400"

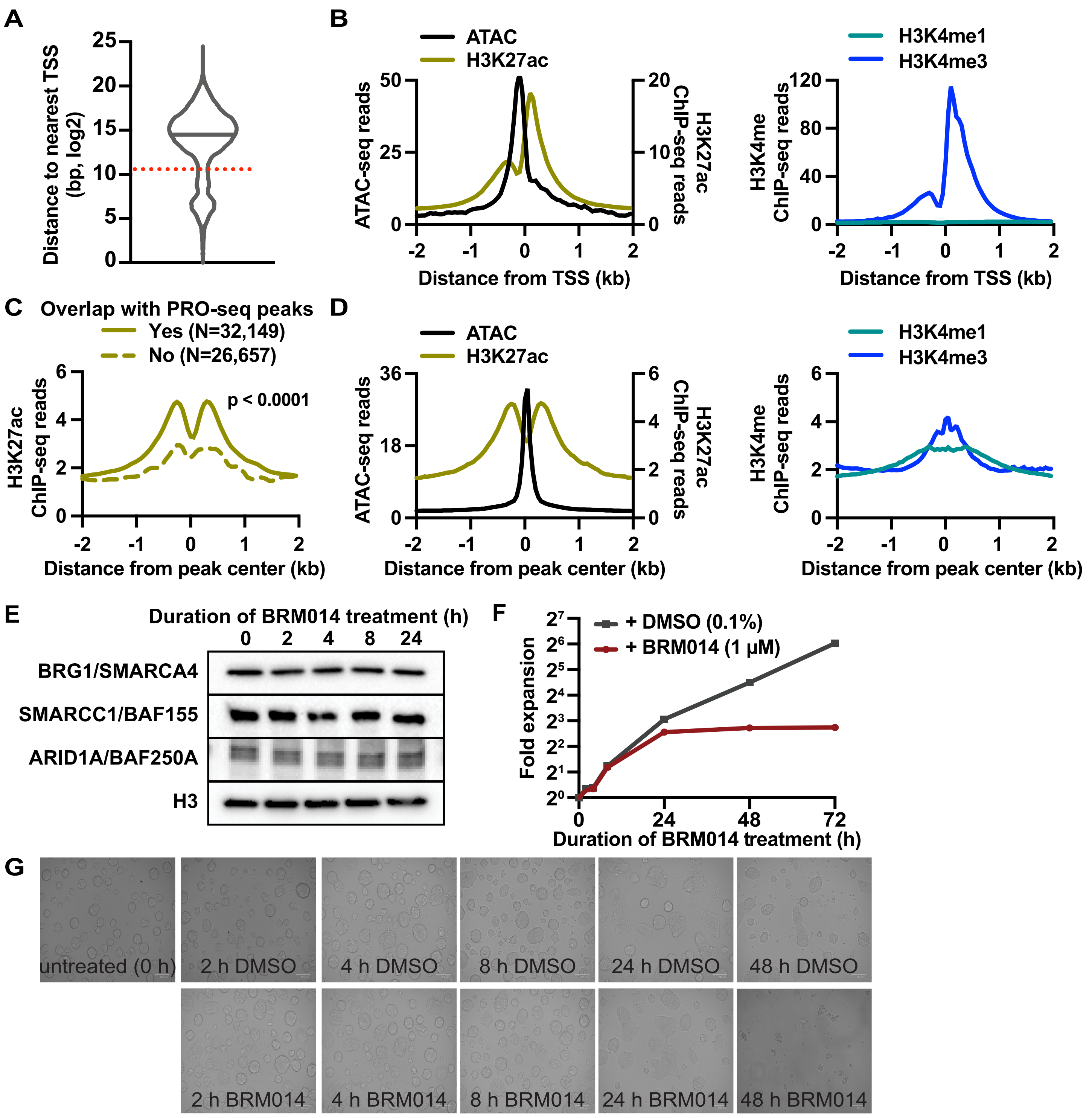

**Figure S1. Promoter and enhancer characteristics and effects of BRM014 treatment in mESCs. Related to Figure 1.**

**(A)** Distribution of distances between all ATAC-seq peaks (n = 83,201) and the nearest active, annotated TSS. Dotted red line indicates 1.5 kb threshold used to separate proximal and distal peaks.

**(B)** Data are shown around proximal peaks, hereafter referred to as promoter peaks (n = 13,536). Average metagene profiles of ATAC-seq and H3K27ac,[60] H3K4me1[61] and H3K4me3[60] ChIP-seq aligned around the associated TSSs, graphed in 50 bp bins.

**(C)** H3K27ac ChIP-seq[60] signal at distal ATAC-seq peaks that do or do not overlap with peaks of PRO-seq signal. Data are graphed in 50 bp bins.

**(D)** For all enhancer peaks (n = 32,149), average profiles of ATAC-seq and H3K27ac,[60] H3K4me1[61] and H3K4me3[60] ChIP-seq are shown, aligned around peak centers, graphed in 50 bp bins.

**(E)** Western blots of cells treated with 1 µM BRM014 for the indicated duration using antibodies against BRG1/SMARCA4, SWI/SNF subunits SMARCC1/BAF155 and ARID1A/BAF250A. Histone H3 is shown as a loading control.
**(F)** Proliferation of mESCs treated with BRM014 or DMSO for the indicated duration.
**(G)** Brightfield images of mESCs treated with 1 µM BRM014 or 0.1% DMSO for the indicated duration.

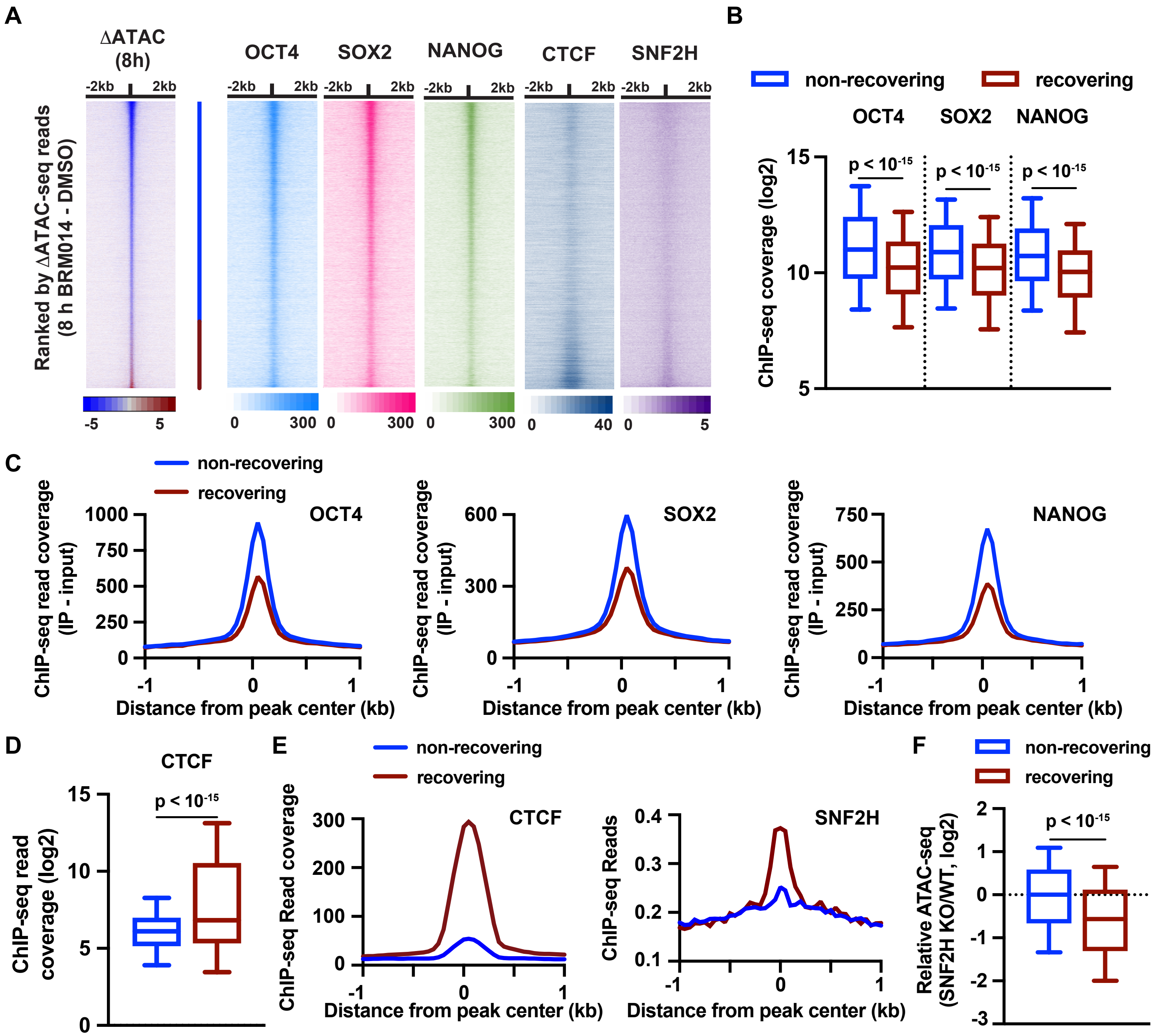

**Figure S2. Enhancers bound by pluripotency-associated pioneer factors require SWI/SNF to retain accessibility, whereas SNF2H opens chromatin at CTCF-associated enhancers. Related to Figure 1.**

**(A)** Heatmap of change in ATAC-seq signal after 8 h BRM014 treatment (ranked as in Figure 1D) across all enhancers is shown alongside heatmaps of ChIP-seq for OCT4,[8] SOX2,[8] NANOG,[8] CTCF[66] and SNF2H,[63] shown in the same rank order. Data are aligned to the enhancer peak center and summed in 50 bp bins. The blue line indicates the positions of the non-recovering enhancers (N=24, 679) and the red line indicates the recovering enhancers (N=7,470).

**(B)** Box plots of ChIP-seq signals for the indicated factors (+/- 300 bp from enhancer center). Whiskers show the 10-90th percentiles and p-values are from Mann-Whitney test.

**(C)** Metagene plots of average ChIP-seq signal (in 50 bp bins) for the indicated TFs, at enhancers separated by whether they recover accessibility following 8 h of BRM014 treatment.

**(D)** Box plots of CTCF[66] ChIP-seq signal (+/- 300 bp from enhancer center). Whiskers show the 10-90th percentiles and p-values are from Mann-Whitney test.

**(E)** Metagene plots of CTCF[66] (left) and SNF2H[63] (right) ChIP-seq signal at enhancers, separated by recovery capacity following BRM014 treatment, graphed in 50 bp bins.

**(F)** For non-recovering and recovering enhancers, relative change in ATAC-seq signal is shown (peak center +/- 300bp) in SNF2H knockout compared to control[4]. Whiskers show the 10-90th percentiles and p-values are from Mann-Whitney test.

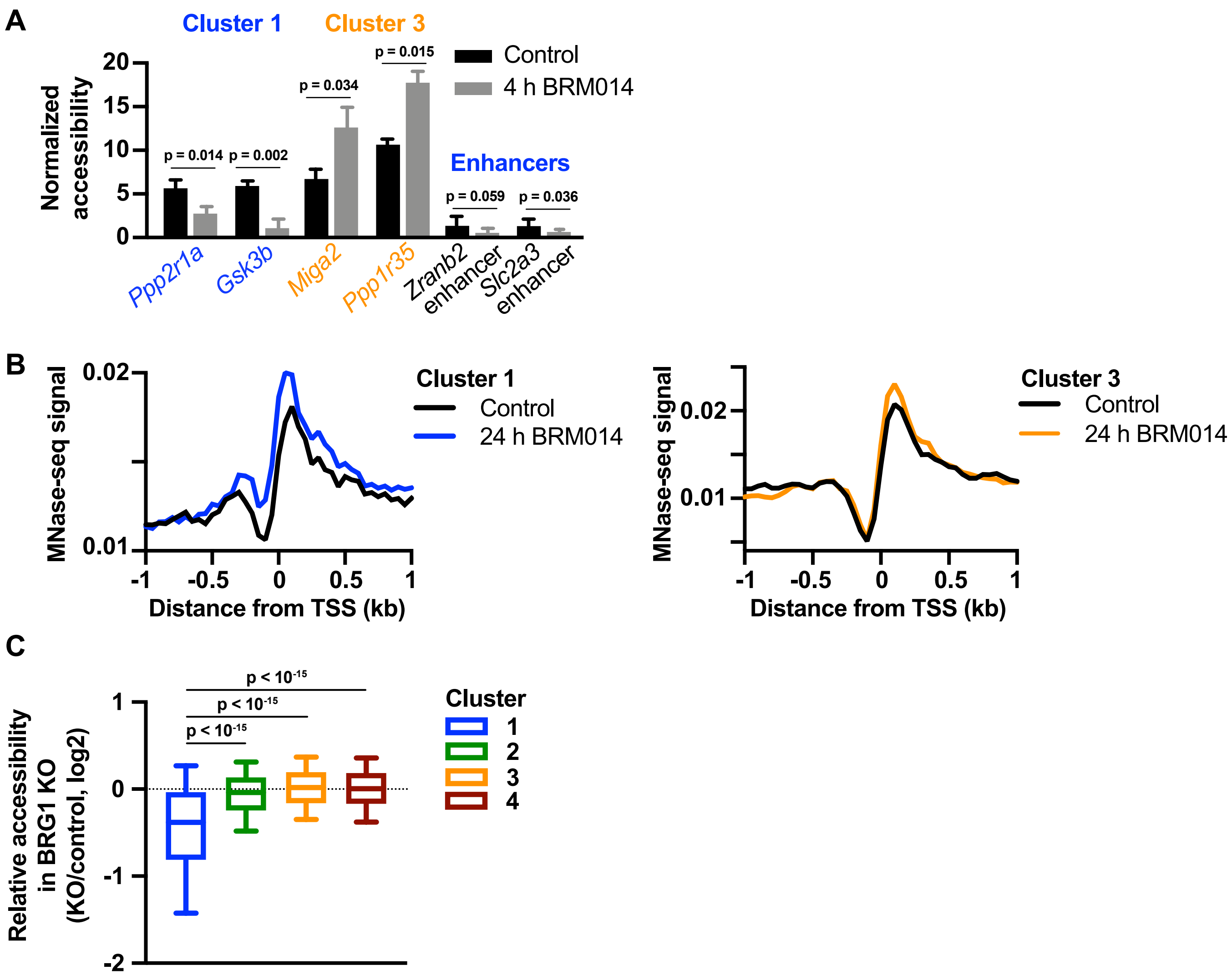

**Figure S3. Cluster 1 promoters show sustained repression of accessibility in the absence of SWI/SNF remodeling. Related to Figure 2.**

**(A)** ATAC-qPCR validation of accessibility changes following 4 h BRM014 treatment at selected promoters without evidence of recovery (Cluster 1, *Ppp2r1a*, *Gsk3b*), promoters with evidence of recovery (Cluster 3, *Miga2*, *Ppp1r35*), and enhancers (*Zranb2* and *Slc2a3* enhancers, neither of which recover). Cq values normalized to background signal of closed chromatin. P-values calculated by Student's t-test.

**(B)** Average MNase-seq signal around Cluster 1 promoters (top panel) and Cluster 3 promoters (bottom panel) in cells subjected to 24 h BRM014 treatment compared to control.[21] Data are graphed in 50 bp bins.

**(C)** Relative accessibility of promoters in each cluster (ATAC-seq signal from -450 to +149 bp relative to TSS) following conditional knockout of BRG1 via 72 h tamoxifen administration in BRG1 fl/fl cells expressing CRE-ER.[8] Whiskers show the 10-90th percentiles and p-values are from Mann-Whitney test.

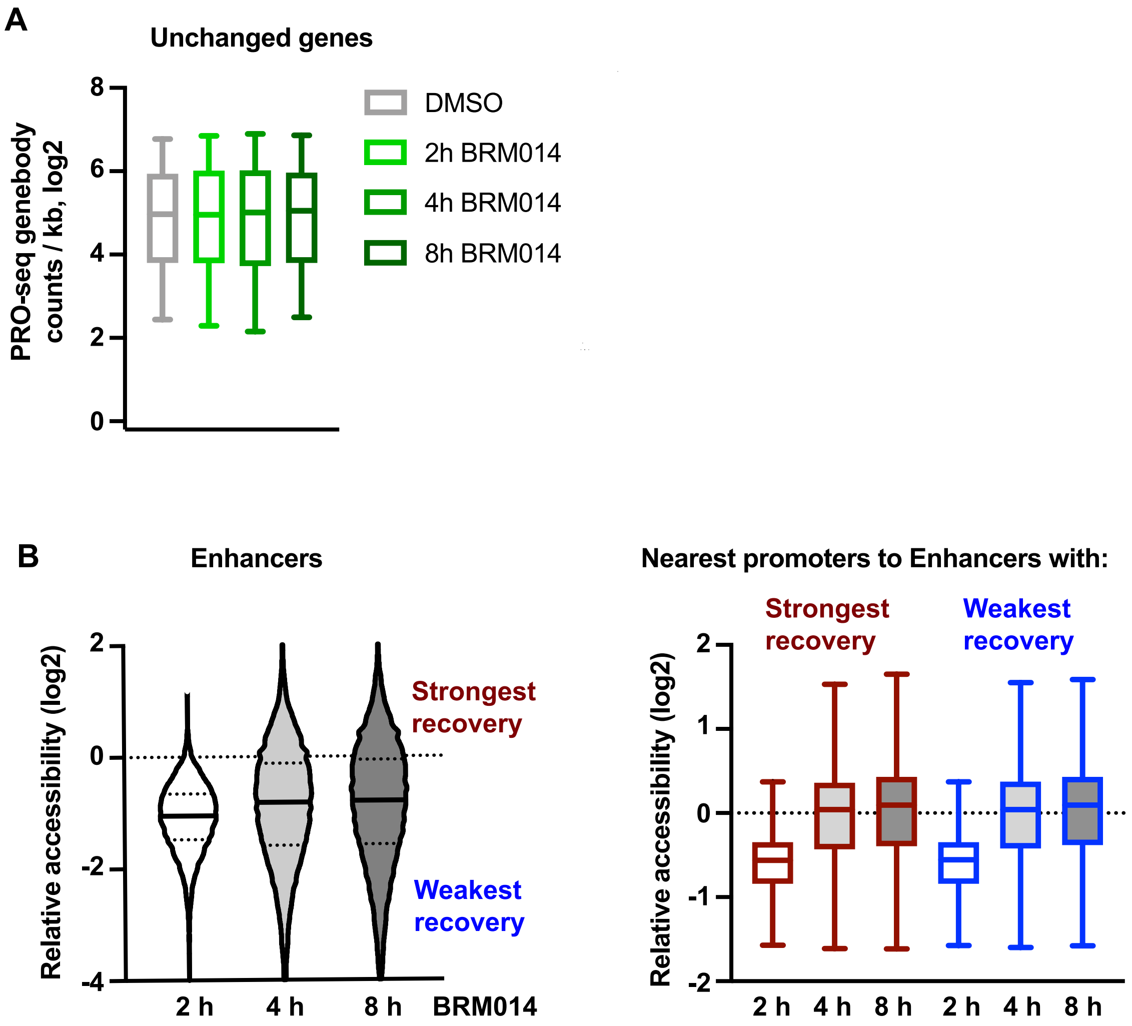

**Figure S4. Enhancer accessibility does not dictate the recovery capacity of nearby genes. Related to Figure 3.**
**(A)** Gene body PRO-seq read density (TSS+250 to TES) at unchanged genes. Whiskers show the 10-90th percentiles. There are no significant differences in PRO-seq reads at 2h, 4h, or 8h BRM014 treatment compared to the DMSO control (Mann Whitney test, p-values all > 0.05).
**(B)** Left: ATAC-seq reads were summed at each enhancer (-300 to +299 bp from the peak center, n = 32,149) over the time course of BRM014 treatment and read counts relative to time matched DMSO samples are shown. As indicated, enhancers were separated into quartiles based on the level of accessibility after 8 h of BRM014. For each enhancer, the nearest gene promoter was identified. Right: The relative ATAC-seq signal is shown at the promoters (-450 to +149 bp relative to TSS) associated with the quartile of enhancers showing the strongest recovery in accessibility (red) and the weakest recovery in accessibility (blue) following 8 h of BRM014 treatment. There is no significant difference in the distribution of relative ATAC-seq signals at promoters associated with the strongest vs. weakest recovering enhancers (Mann-Whitney test, p-value > 0.05). Whiskers show the 10-90th percentiles.

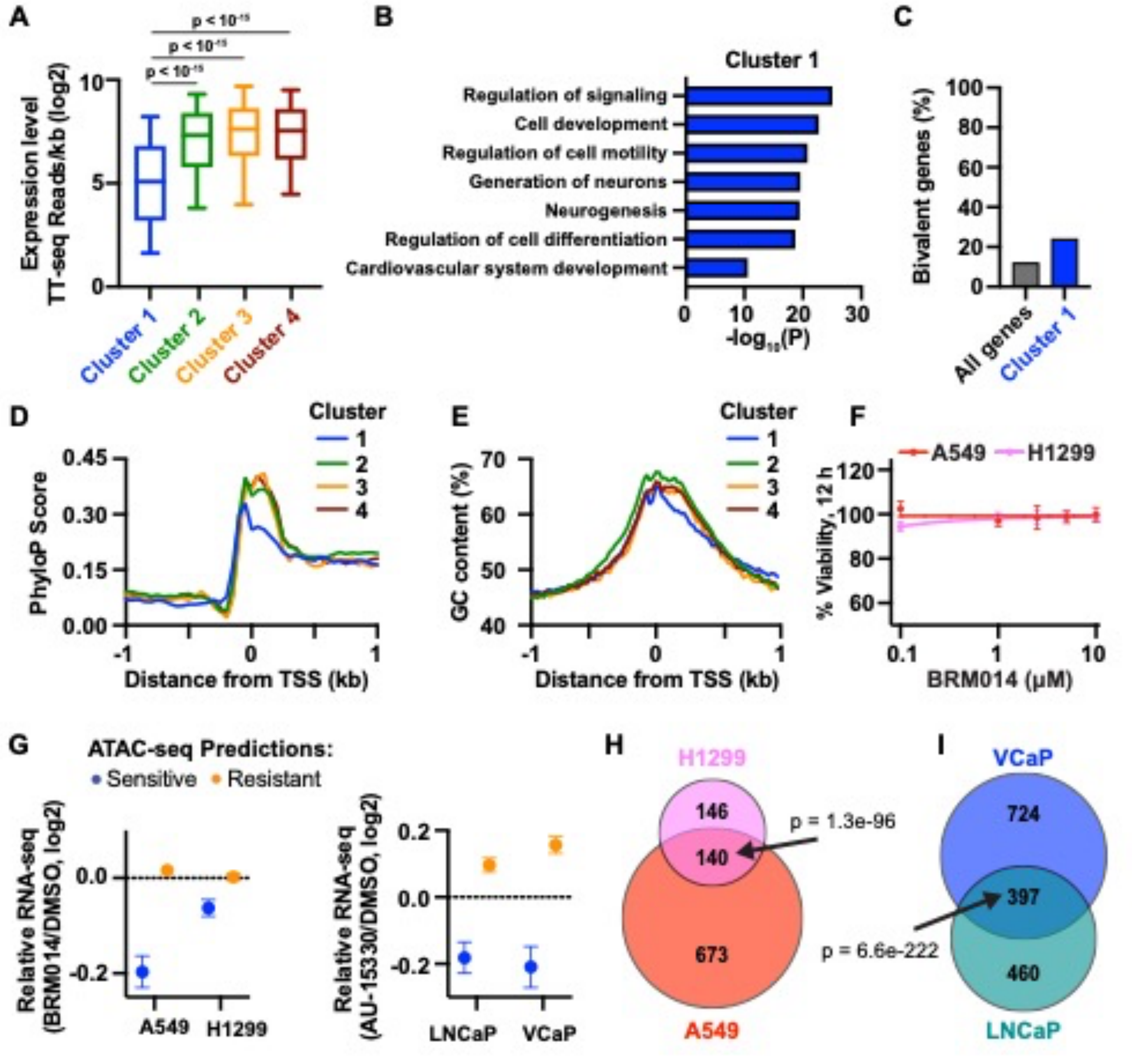

**Figure S5. Cluster 1 genes exhibit unique characteristics that are conserved in human cancer cells. Related to Figures 4 and 5.**

**(A)** TT-seq[60] signal at genes by promoter cluster. Whiskers show the 10-90th percentiles and p-values are from Mann-Whitney test.

**(B)** Select GO terms enriched in cluster 1 genes, using cluster 2 genes as background.

**(C)** Percentage of genes classified as bivalent.[91]

**(D)** Average PhyloP conservation score[62] for promoters by cluster, graphed in 50 bp bins. A higher PhyloP score indicates a higher level of evolutionary conservation, as determined across placental species.

**(E)** Average GC content[62] at promoters by cluster, graphed in 25 bp bins.

**(F)** Drug dose-response curves of BRM014 treated A549 or H1299 cell lines after 12 hours of treatment. Viable cells counted using CellTiter-Glo. Error bars represent SEM of six technical replicates.

**(G)** Mean expression changes at genes predicted to be sensitive to SWI/SNF perturbation using ATAC-seq. The average log2 fold-change in RNA-seq following SWI/SNF inhibition by BRM014 (A549 and H1299 cells) or degradation by AU-15330[25] (LNCaP and VCaP cells) is shown. Error bars represent SEM. See Methods for data sources and number of genes in each group.

**(H-I)** Overlap between genes downregulated (Fold-change > 1.5 and P adj < 0.001) following BRM014 treatment in H1299 and A549 cells (H) or AU-15330 treatment in VCaP and LNCaP cells[25] (I). P-values for the overlap of gene lists were calculated using the hypergeometric distribution with the phyper function in R.

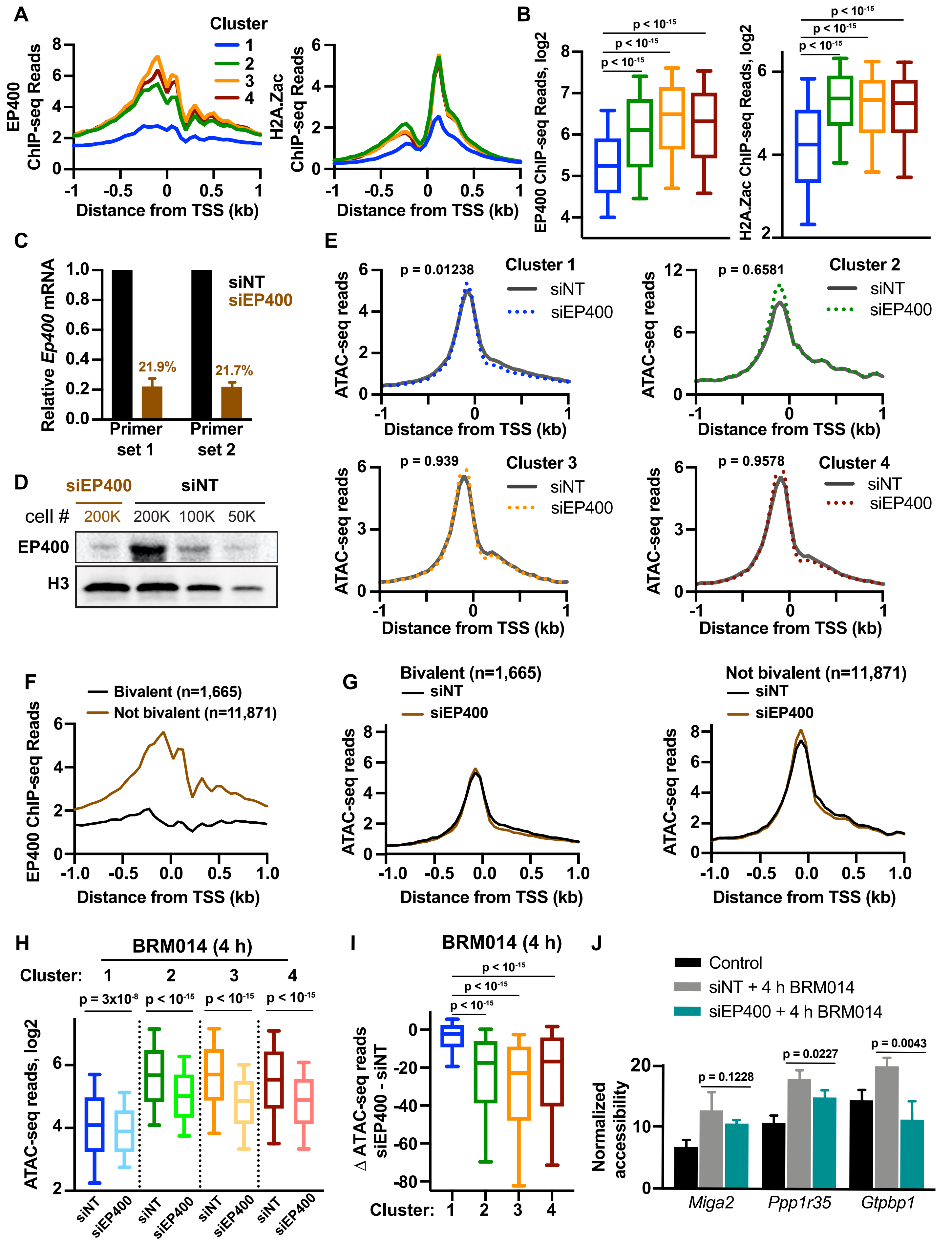
A
Cluster
1
2
3
4
EP400 ChIP-seq Reads
H2A.Zac ChIP-seq Reads
Distance from TSS (kb)
B
EP400 ChIP-seq Reads, log2
H2A.Zac ChIP-seq Reads, log2
p < 10-15
C
Relative Ep400 mRNA
siNT
siEP400
21.9%
21.7%
Primer set 1
Primer set 2
D
siEP400
siNT
cell #
200K
200K
100K
50K
EP400
H3
E
Cluster 1
Cluster 2
Cluster 3
Cluster 4
p = 0.01238
p = 0.6581
p = 0.939
p = 0.9578
ATAC-seq reads
F
Bivalent (n=1,665)
Not bivalent (n=11,871)
EP400 ChIP-seq Reads
G
Bivalent (n=1,665)
Not bivalent (n=11,871)
H
BRM014 (4 h)
Cluster:
p = 3x10-8
ATAC-seq reads, log2
I
BRM014 (4 h)
Δ ATAC-seq reads siEP400 - siNT
J
Control
siNT + 4 h BRM014
siEP400 + 4 h BRM014
Normalized accessibility
p = 0.1228
p = 0.0227
p = 0.0043
Miga2
Ppp1r35
Gtpbp1

**Figure S6. siRNA-mediated knockdown of EP400 for 72 h substantially reduces levels of EP400 mRNA and protein and impairs promoter recovery following BRM014 treatment. Related to Figure 6.**

**(A)** Average EP400[3] and H2A.Zac[65] ChIP-seq reads by promoter cluster, graphed in 50 bp bins.

**(B)** For promoters by cluster, box plots of EP400[3] and H2A.Zac[65] ChIP-seq reads per gene (TSS +/- 500bp). Whiskers show the 10-90th percentiles and p-values are from Mann-Whitney test.

**(C)** RT-qPCR analysis of *Ep400* mRNA levels following 72 h treatment with siNT or siEP400. Two primer pairs targeting *Ep400* were used. Data was normalized for each primer pair by levels detected in siNT treated cells.

**(D)** Western blot analysis of EP400 protein levels following 72 h treatment with siNT or siEP400. Histone H3 is shown as a loading control.

**(E)** For promoters by cluster, average ATAC-seq signal in siNT treated and siEP400 cells, graphed in 50 bp bins. To assess differences between conditions, ATAC-seq signal was summed (-450 to +149 bp relative to TSS) in siNT treated and siEP400 cells. P-values calculated by Wilcoxon signed ranks test.

**(F)** Average EP400[3] ChIP-seq reads for genes classified as bivalent[91] vs. those not classified as bivalent, graphed in 50 bp bins.

**(G)** For promoters classified as bivalent[91] (left) or not (right), average ATAC-seq signal in siNT treated and siEP400 cells, graphed in 50 bp bins.

**(H)** For promoters by cluster, box plots of ATAC-seq signal (-450 to +149 bp relative to TSS) in siNT and siEP400 cells following 4 h BRM014 treatment. Whiskers show the 10-90th percentiles and P-values are from Mann-Whitney test.

**(I)** For promoters by cluster, boxplots of the difference in ATAC-seq signal (-450 to +149 bp relative to TSS) in BRM014 treated cells (siEP400 minus siNT). Whiskers show the 10-90th percentiles and p-values are from Mann-Whitney test.

**(J)** ATAC-qPCR validation of accessibility changes following 4 h BRM014 treatment in siNT and siEP400 cells at selected promoters with evidence of recovery (*Miga2*, *Ppp1r35*, *Gtpbp1*). qPCR values were normalized to background signal of closed chromatin. P-values calculated by Student's t-test.

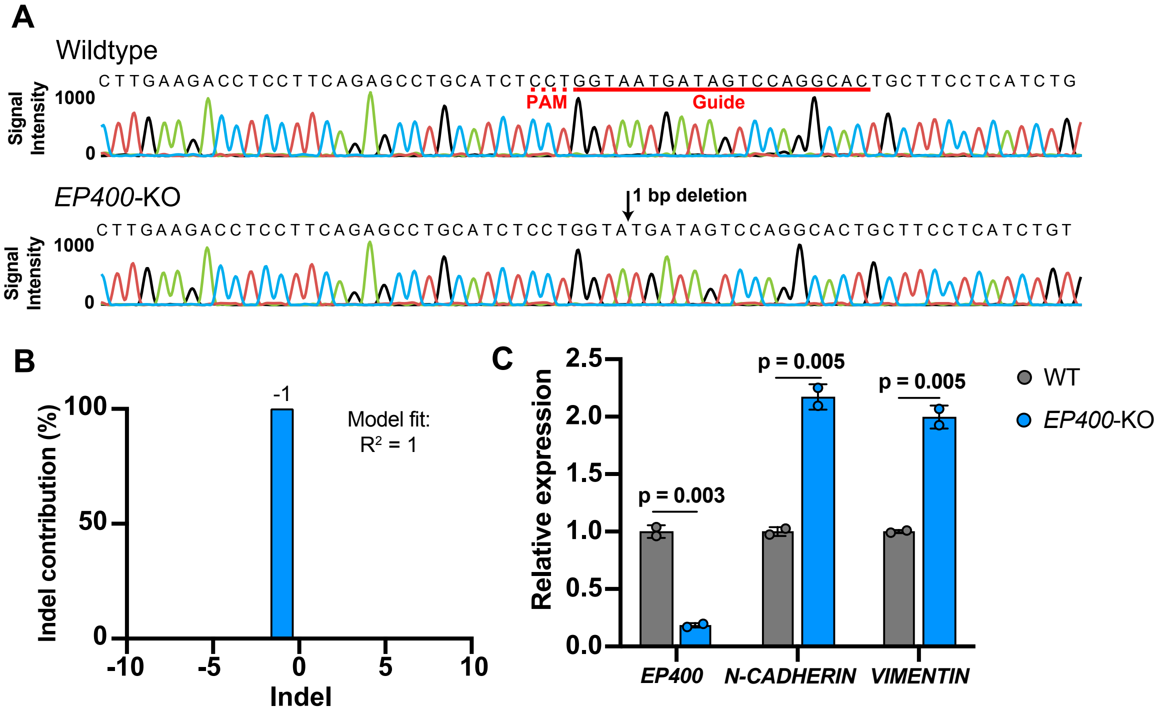

**Figure S7. Characterization of *EP400*-KO A549 cell line. Related to Figure 7.**

**(A)** Sanger sequencing traces spanning the CRISPR guide cut site (in the coding sequence of *EP400* exon 2) from wildtype and *EP400*-KO cell lines were compared using Inference of CRISPR Edits (ICE).[87] Raw sequencing traces shown with CRISPR guide and 1 bp deletion in KO indicated.

**(B)** Indel contributions shown for the *EP400*-KO cell line, indicating a homozygous 1 bp deletion.

**(C)** *EP400*, *N-CADHERIN* and *VIMENTIN* transcript levels were assessed by RT-qPCR in parental (wildtype) and *EP400*-KO cells (n = 2 for each condition). *EP400* expression was normalized to *ActB* and then to the average expression in wildtype cells. Primers for *EP400* anneal to exons 29 and 30. Average expression per genotype plotted, with individual values shown as circles and error bars representing the SEM. P-values calculated by Student's t-test.

### Supplementary Tables

#### Table S1. Oligonucleotide sequences used in this study. Related to STAR Methods.

**Table S1. Oligonucleotide sequences used in this study. Related to STAR Methods.**

| **RT-qPCR primers** | **Fwd** | **Rev** |
|---|---|---|
| human EP400 | ATGAAAATGCCAGCAGTGAGC | TGTGAGTGGAGAGAATGGCAC |
| human VIMENTIN | AAAGTGTGGCTGCCAAGAAC | AGCCTCAGAGAGGTCAGCAA |
| human N-CADHERIN | CCACAATCCTGTCCACATCT | TTCGGGTAATCCTCCCAAAT |
| human ActB | CCGCCGCCAGCTCAC | CCACGATGGAGGGGAAGAC |
| mouse EP400 #1 | GCAGATGCAGACATCTCAGC | GTTGGTAGCAGAAGAGGGTC |
| mouse EP400 #2 | GGACTGGTTGGCCAAGCTAT | CTGCACAGTTTTCCCAAGCC |
| **ATAC-qPCR primers** | **Fwd** | **Rev** |
| Ppp2r1a | TCTGGCCGCTCTATTTCTTC | GATTTGCCTTGCTTGGTTTC |
| Gsk3b | TCGCCGTCCTTAACTCTTGA | GTCCTGTGGCTGGCTAGAGA |
| Miga2 | ACCAGATCAGCAGGGCTAGA | CTCATCCCACCTTGGACACC |
| Ppp1r35 | AAGGCGAAGTTAGCGTGTGA | CTCCTGGACGTCATTCCCAC |
| Zranb2-enhancer | CACATCTCCCTGCAACACAG | CAAGTCTCCACCAACCATTTC |
| Slc2a3-enhancer | GATCCCATGCAGGAAATGAC | GGAAGGGAACTCAGGGAAAG |
| Gtpbp1 | CTGTGTTGGCCCCACTAGAG | TACCAATCGTCGCAGCTCTC |
| Background 1 | GGACTGGGAGGGAGGAAGT | CATTCTGTCTGCTGGCTTCA |
| Background 2 | CACCACTCCTCAAGGGATGT | CCTCGGTGGATGTCTGTACC |
| Background 3 | TTCATGTCCTGGTTGGTTTG | CCTCCCTTGACTTCCCTTTC |
| **Genotyping primers** | **Fwd** | **Rev** |
| EP400 | TGTGTTTCAGGATGGGTCAG | CTGAGTTCTCTGAACCTGAGATG |
| **CRISPR** | **sgRNA protospacer** | **crRNA sequence** |
| EP400 exon 2 | GTGCCTGGACTATCATTACC | GUGCCUGGACUAUCAUUACCGUUUUAGAGCUAUGCU |
